# Splicing dysregulation in glioblastoma alters the function of cell migration-related genes

**DOI:** 10.1101/2024.10.05.616792

**Authors:** Minu Seong, Pedro Bak-Gordon, Zhaoqi Liu, Peter D. Canoll, James L. Manley

## Abstract

Glioblastoma (GBM) has a poor prognosis with a high recurrence and low survival rate. Previous RNA-seq analyses have revealed that alternative splicing (AS) plays a role in GBM progression. Here, we present a novel AS analysis method (Semi-Q) and describe its use to identify GBM-specific AS events. We analyzed RNA-seq data from normal brain (NB), normal human astrocytes (NHAs) and GBM samples, and found that comparison between NHA and GBM was especially informative. Importantly, this analysis revealed that genes encoding cell migration-related proteins, including filamins (FLNs) and actinins (ACTNs), were among those most affected by differential AS. Functional assays revealed that dysregulated AS of *FLNA, B* and *C* transcripts produced protein isoforms that not only altered transcription of cell proliferation-related genes but also led to enhanced cell migration, resistance to cell death and/or mitochondrial respiratory function, while a dysregulated AS isoform of *ACTN4* enhanced cell migration. Together, our results indicate that cell migration and actin cytoskeleton-related genes are differentially regulated by AS in GBM, supporting a role for AS in facilitating tumor growth and invasiveness.

## Introduction

GBM is one of the most infiltrative and invasive cancer types, with a five-year survival rate of only 6.8%. Treatment for GBM is limited to surgical removal, temozolomide chemotherapy and radiotherapy, and 90% of patients experience recurrence (Preusser et al. 2011; Geraldo et al. 2019; Tan et al. 2020). Drug development toward new treatment targets continues, but clinically available drugs are still insufficient (Nguyen et al. 2018; Chelliah et al. 2021; Poonan et al. 2021). Recently, large-scale RNA-seq analyses revealed that differential AS in GBM is a prominent molecular feature, and that such AS differences could provide promising therapeutic targets (Babenko et al. 2017; Zhao et al. 2017; Li et al. 2019; Wang et al. 2021). Although the results vary depending on the type and number of samples and the programs used to analyze RNA-seq data, several prognostic markers, splicing regulators, glioma-specific neoantigens and signaling pathways have been identified.

Current computational tools for AS analysis mainly fall into two categories: Isoform-based (e.g., Cufflinks and DiffSplice) and local event-based (e.g., JUM, rMATS and MAJIQ) methods (Wang and Rio 2018; Liu and Rabadan 2021) methods. The isoform-based programs build full-length transcripts of different isoforms and then compare them with each other. However, reconstruction of full-length transcripts can be difficult and error-prone if the transcript is too long or if there are numerous isoforms. The event-based programs use specific exon and junction information and tend to be widely used because many studies focus on local AS events. One of the most challenging steps of AS analysis programs is isoform-level quantification. It is possible, however, to convert FPKM (Fragments Per Kilobase of transcript per Million mapped reads) values into FPKM ratios and use the latter for AS analysis. Such an approach has the potential to simplify the quantification process greatly, but is a semi-quantitative method, and has not yet been shown to be entirely reliable.

A number of AS events are enriched in the brain and, as mentioned above, may be dysregulated in GBM. For example, microexons are short exons, typically defined as <30nt, that play an important role in the development and maintenance of neuronal function (Volfovsky et al. 2003; Li et al. 2015; Ustianenko et al. 2017). The discovery and early studies of microexons were limited to neuronal cells, but muscle- and microglia-specific microexons have been identified and their functions studied (Lee et al. 2022). It has also been suggested that a possible subset of glia-specific microexons might be skipped in glioma (Reixachs-Solé et al. 2020). Therefore, cell subtype-specific microexon inclusion can be used as an indicator of cell-type specificity for AS analyses, as well as a possible target of splicing dysregulation in glioma.

Another factor influencing analysis of cancer-specific splicing derives from the fact that non-tumor cells closely communicate and cooperate with tumor cells. Tumor purity is highly associated with clinical and molecular features, as non-tumor cells play crucial roles in tumor proliferation, invasion and angiogenesis (Rape et al. 2014; Yadav and De 2015). In glioma, tumor impurity arises primarily from the presence of immune cells such as macrophages and microglia, which can constitute up to 30% of tumor mass (Komohara et al. 2008), and low purity is correlated with clinical malignancy and reduced survival time (Zhang et al. 2017). Thus glioma biopsies often reveal the presence of immune cells (Quail et al. 2016; Quail and Joyce 2017). In addition to contributing to the microenvironment of the tumor, the presence of immune cells in tumor biopsies can complicate molecular characterization of GBM.

In this study, we have characterized global changes in GBM-specific AS and determined the functional significance of several key AS events. To facilitate this analysis, we first developed and validated a new computational method for evaluating AS changes, called Semi-Quantitative RNA-seq analysis (Semi-Q). Using this method, we observed significant microexon exclusion in GBM compared to normal brain (NB) samples. However, we concluded this was due to the presence of neuronal cells in the NB samples, and that microexons are not in fact specifically excluded by AS in GBM. This suggested that NB may thus not be optimal for AS comparison with GBM, and we therefore used normal human astrocytes (NHAs) instead of NB for comparison. Strikingly, we found that multiple cell migration-related genes were amongst the most differentially regulated by AS between NHA and GBM. After verifying a number of these by RT-PCR with postmortem GBM brains and cultured cells, we examined the functional significance of several, specifically actinins (*ACTN*s) and filamins (*FLN*s). Notably, mutually exclusive exon 8B inclusion in *ACTN4* enhanced cell migration, while skipping of a “hinge” exon in *FLN A, B* and *C* produced isoforms that both altered transcriptional activity, thereby increasing expression of pro-growth genes, while also directly enhancing several functions related to tumor cell growth. Together, our results support a role for AS in facilitating tumor growth and invasiveness in GBM.

## Results

### Comparative RNA-seq analysis reveals differences between NB and GBM

To gain insights into the extent and role of dysregulated AS in GBM, we first downloaded RNA-seq data from 90 NB samples in the Genotype-Tissue Expression (GTEx) Project, and from 155 GBM samples in The Cancer Genome Atlas (TCGA) (Supplemental Table S1). To ensure an accurate comparison between these samples, we selected RNA-seq datasets of high quality using Gene Body analysis, which gave rise to 47 NB and 143 GBM samples (Supplemental Fig. S1 and Supplemental Table S2; see Materials and Methods for details). To focus on high-grade gliomas, we sorted the 143 GBM samples based on subtype. While high-grade gliomas were initially classified into four molecular subtypes, classical, mesenchymal, proneural and neural (Verhaak et al. 2010), the neural subtype was found to arise from contamination of neuronal tissue, and is no longer considered GBM (Sturm et al. 2012; Gill et al. 2014; Behnan et al. 2017). Among our 143 GBM samples, 35 were classical, 47 were mesenchymal and 26 were proneural, leaving 108 GBM samples in the three GBM molecular subtypes that were high-grade gliomas. Further quality control analyses eliminated one additional GBM and one NB sample (see Materials and Methods for more details of the classification).

We next wished to use these high-quality samples to investigate global changes in AS that occur in GBM. To this end, the NB and GBM samples were analyzed in two different ways. One involved a standard Cufflinks-Cuffquant-Cuffdiff process (Trapnell et al. 2012), and the other the Semi-Q method that we developed (see Materials and methods and Supplemental Fig. S2). We decided to focus on coding cassette exons (CCEs), because inclusion/exclusion of such exons is the most common AS mechanism and is a major contributor to the functional diversity of proteins (Wang et al. 2017). 1078 CCEs from 655 genes were obtained using the Cuffdiff method (Supplemental Table S3), and 952 CCEs from 645 genes were identified using Semi-Q (Supplemental Fig. S2D and Supplemental Table S3). For completeness, we also analyzed CCEs using the event-based method rMATS and identified 1333 CCEs from 880 genes (Shen et al. 2014) (Supplemental Table S3).

Neither NB nor GBM tissues consist of pure cell populations. NB samples are a mixture of glia and neurons, while GBM typically consists of gliomas containing variable fractions of neurons, microglia, macrophages and non-neoplastic glia, including astrocytes and oligodendrocytes (Gill et al. 2014; Quail et al. 2016; Quail and Joyce 2017; DeCordova et al. 2020; Al-Dalahmah et al. 2023). To determine whether this would be apparent in our analyses, we performed Gene Ontology (GO) analysis with the NB and GBM CCE-containing gene groups using results generated by Semi-Q, Cuffdiff and rMATS. The GO results indicate that neuronal function-related terms were pronounced in the NB-specific CCE-containing gene group in all three analyses. Notably, however, immune function-related terms were pronounced in the GBM-specific CCE-containing genes, but only with the Semi-Q results (Supplemental Fig. S3A, Supplemental Table S4). Why this was the case is not clear but given that this result is consistent with the known presence of immune cells in GBM (see above), we conclude that Semi-Q provides an accurate representation of the transcripts present in these samples.

Amongst the differentially spliced CCEs, we found that microexons appeared to be frequently excluded in GBM samples relative to the NB samples. The length distribution of CCEs generated by all three of the above methods was analyzed, and the results suggested that microexons were often excluded in GBM, although the percentages varied (Fig. 1A). However, microexons are known to be specifically included primarily in neurons, not in glia (Irimia et al. 2014; Gonatopoulos-Pournatzis and Blencowe 2020). Thus, neuronal cells in the NB samples may have been the main source of microexon inclusion, giving rise to the apparent relative exclusion in GBM. To explore this hypothesis, microexon inclusion status was analyzed by comparing NB samples, NHA cells and A172 GBM-derived cells, by RT-PCR. As shown in Figure 1B, microexons were excluded in both the NHA and A172 cells, but were included in NB. We conclude that the microexon inclusion we detected in NB samples indeed arose primarily from neuronal cells and not glia, consistent with the studies referenced above showing that microexon inclusion is neuron-specific. Thus, while microexons are excluded in GBM, this does not reflect a change in AS during GBM tumorigenesis (see also below). This finding suggests that a different comparison than NB vs GBM could provide a more accurate depiction of GBM-specific changes in AS.

**Figure 1.**
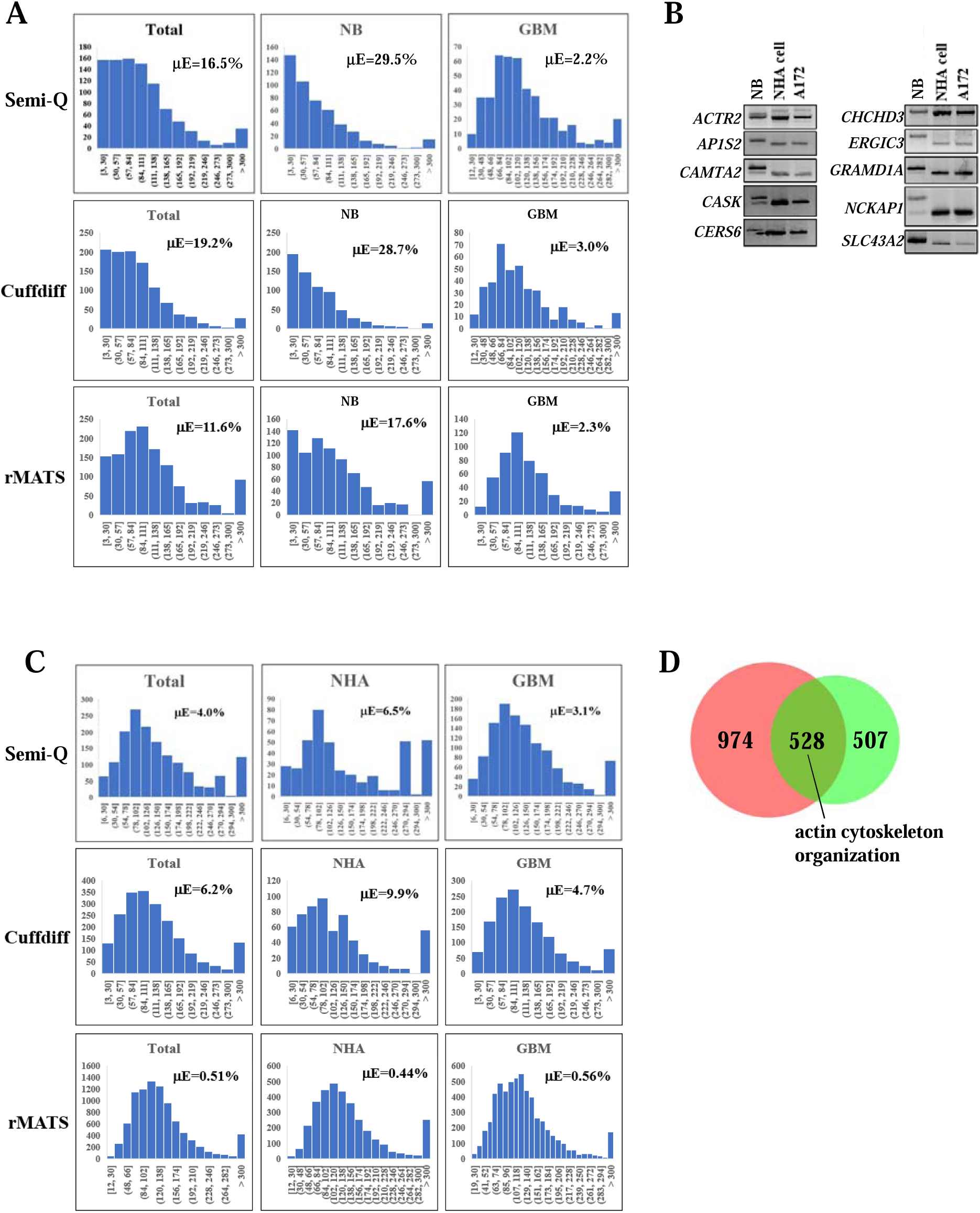
Normal brain biopsies constitute a mixture of cell types, which presents challenges for comparative RNA-seq analysis with GBM. A. The length distribution of CCEs in NB and GBM is represented by histograms. Three different programs (Semi-Q, Cuffdiff and rMATS) were used, and three different groups of CCEs (total CCEs, NB-specific CCEs, and GBM-specific CCEs) were analyzed. The horizontal axis represents the interval of exon length (in nt), and the vertical axis represents the number of exons. The percentage of microexons (µE) out of total exons is indicated. B. The indicated µE-containing genes (vertical axis) were selected and µE inclusion in NB, NHA and A172 cells was analyzed by RT-PCR. NB, normal brain, NHA, normal human astrocytes, and A172, a glioma cell line. C. The length distribution of CCEs in NHAs and GBM is represented by histograms. Analysis was identical to that shown in Figure 1A. D. CCE- containing genes that displayed differences in AS between NHA and GBM by Semi-Q (green circle) and Cuffdiff (red circle) were compared and overlapping genes are indicated (olive green). The numbers of genes in each category are indicated. GO analysis of the overlapping genes revealed genes involved in “actin cytoskeleton organization.”

### Comparative RNA-seq analysis between NHA and GBM samples reveals extensive AS changes

In light of the above, we reasoned that a comparison between NHAs and GBM might be more reflective of physiological differences in AS than one between NB and GBM. To investigate this, we downloaded from the Sequence Read Archive database eleven NHA RNA-seq datasets that had been obtained from NHAs isolated from individual human brains. These datasets were processed as described in Materials and Methods (Supplemental Table S1), and after discarding one because of poor quality, the ten remaining NHA samples were compared with the 107 GBM samples in the same way as were the GBM and NB samples. 2093 CCEs from 1502 genes were identified using the Cufflinks-Cuffquant-Cuffdiff process, and 1608 CCEs were obtained from 1035 genes using Semi-Q (Supplemental Table S5). We also identified 9181 CCEs from 3678 genes using rMATS (Supplemental Table S5).

We first analyzed the results from these comparisons for features described above, i.e., microexon inclusion. In contrast to our findings generated by comparing NB with GBM, which suggested that GBM specifically excludes microexons, the length distribution of CCEs revealed that relative to the NHA samples, changes in microexon exclusion in GBM were minimal, only ∼2 fold as opposed to the >10 fold observed when comparing NB and GBM (Fig. 1C). As noted above, microexons are included specifically in neurons and not in glia, which explains the discrepancy between these comparisons. Another notable feature of our analysis was that microexons in NHAs accounted for about 6-10% of total CCEs using the Semi-Q and Cuffdiff methods but the fraction detected by rMATS was much lower, <0.5% (Fig. 1C). The former results are more in keeping with previous studies (e.g., Reixachs-Solé et al. 2020), and we are unsure why microexons were underrepresented by rMATS.

For further analysis, we decided to concentrate on the CCE-containing genes that displayed AS differences between NHAs and GBM by both Cuffdiff and Semi-Q (528, see Fig. 1D). Interestingly, GO analysis revealed functions closely related to cell migration, and we selected genes from among the 40 in the group ‘actin cytoskeleton organization’ for further detailed study (Table 1 and 2, yellow highlighted). We chose this group in light of the invasiveness and aggressiveness of GBM, and the likelihood that alterations in the cytoskeleton are important for these properties (Keller et al. 2022). We first verified AS changes of select transcripts from this group by RT-PCR. For this analysis, we chose several genes with potential cancer relevance, specifically *ACTN2/4, CORO7, FLNB/C* and *GSN/VILL,* and used RNA isolated from both NHA and A172 and U87 GBM cells, as well as from post-mortem human NB and low- and high- grade glioma samples (Fig. 2A). As described below, in all cases the results largely confirm the differential AS detected from our Cuffdiff/Semi-Q analyses, and strikingly the AS changes in GBM all suggest production of protein isoforms that have the potential to enhance cell proliferation.

**Figure 2.**
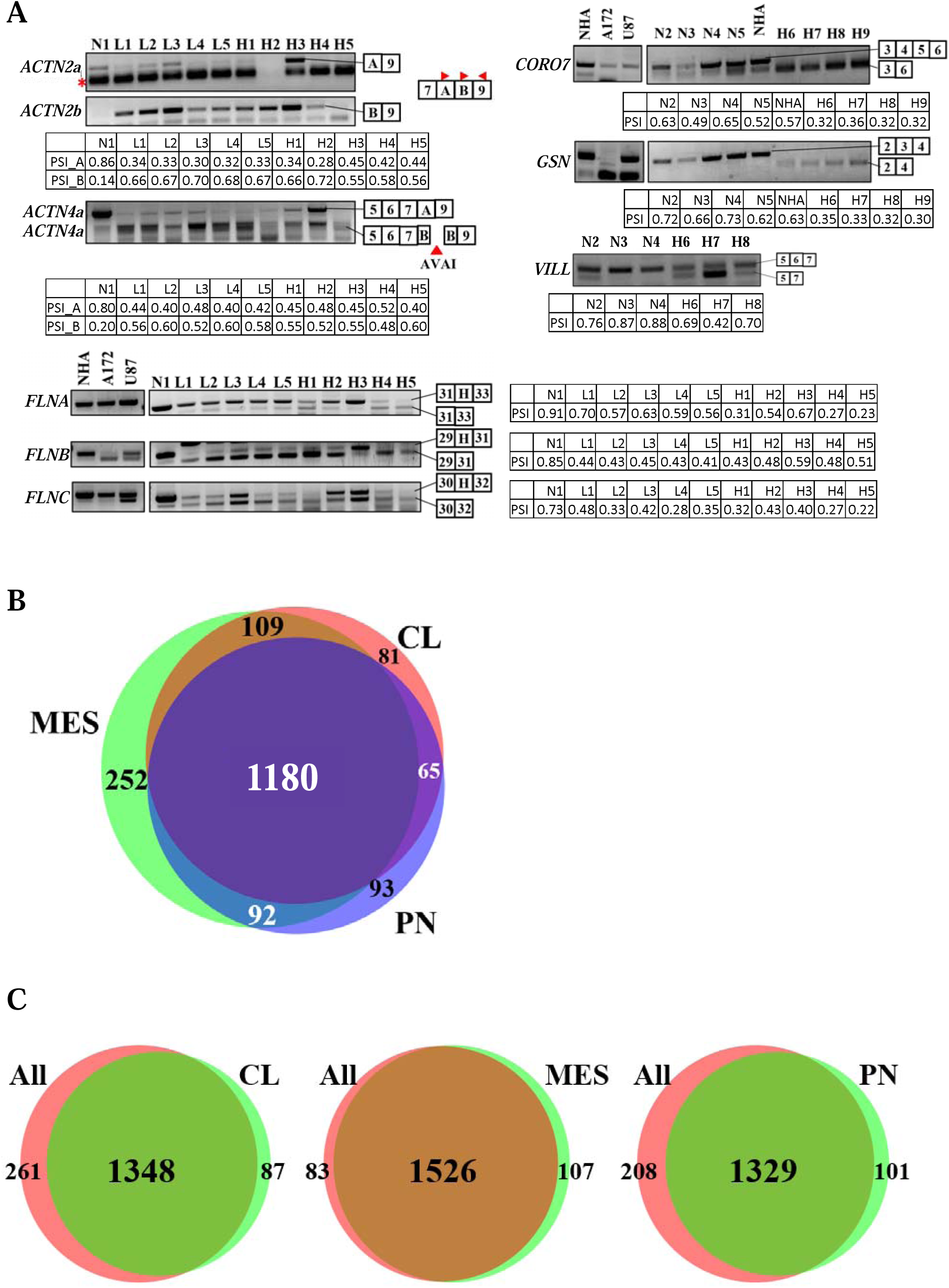
Confirmation and characterization of AS changes detected by RNA-seq comparison of NHAs with GBM. A. AS of CCEs in select genes (indicated on the left) from the “actin cytoskeleton group” identified in Fig. 1D was verified by RT-PCR, using appropriate primers. For *ACTN2*, exon 8A-specific or exon 8B-specific forward primers and exon 9-specific reverse primers were used. For *ACTN4*, exon 5-specific forward primer and exon 9-specific reverse primer were used, and PCR fragments underwent AVAI restriction enzyme digestion after PCR reaction. For filamins, nested hinge exon-specific primer sets were used. For *CORO7, GSN* and *VILL*, nested CCE exon specific primer sets were used. N1-N5, NB samples; L1-L5, low-grade glioma samples; H1-H9, high-grade glioma samples. Exons included are indicated on the right. Asterisk indicates a nonspecific band. PSI (percent spliced in) values are shown in the adjacent tables. B. NHA and each of the three GBM subtypes were compared, and subtype-specific CCEs were identified. The three groups of subtype-specific CCEs were compared with one another. MES, mesenchymal; PN, proneural; CL, classical. C. Comparison between dysregulated CCEs in each of the three subtypes (MES, PN and CL) and CCEs from the comparison between NHA and all GBM samples (All).

**Table 1.** GO analysis using the 528 genes in Figure 1D revealed that cell migration related genes are dysregulated by AS in GBM (yellow highlighted).

**Table 2.** List of genes in the actin/cytoskeleton organization group. Genes validated by RT-PCR are highlighted in yellow.

### Alternative splicing of multiple transcripts encoding cytoskeletal regulators is dysregulated in GBM

Actinins are actin cross-linking proteins that function to enhance stress fiber formation and cell migration. Notably, exons 8A and 8B of *ACTN4* are mutually exclusive, and inclusion of exon 8B is frequently found among small-cell lung cancer and neuroendocrine pulmonary tumor patients, and is correlated with a low survival rate (Miyanaga et al. 2013; Honda 2015; Okamoto et al. 2015). Exon 8B inclusion is also known to enhance stress fiber formation (Honda et al. 2004). While less studied, *ACTN2* has a similar gene structure as *ACTN4* (Lek et al. 2010; de Almeida Ribeiro et al. 2014). Notably, the AS pattern of mutually exclusive exons in *ACTN2/4* transcripts is similar to that of Pyruvate Kinase M, which is also dysregulated in GBM (David et al. 2010). RT-PCR results confirmed that exon 8B of both *ACTN2* and *ACTN4* was largely excluded in the NB sample, but included in the low- and high- grade gliomas, consistent with the Cuffdiff/Semi-Q analyses, which both revealed enhanced exon 8 exclusion in GBM.

Filamins are also actin-binding proteins. They (FLN A, B and C) function in part to bridge integrins to filamentous actin (Savoy and Ghosh 2013). After the integrin signal is transduced, FLNA (and likely B and C) functions as a signal adaptor, and is rapidly cleaved in a “hinge” region. The truncated N- and C-termini are translocated to the nucleus and interact with several transcription factors to suppress tumor growth and inhibit metastasis (Bedolla et al. 2009; Deng et al. 2012; Mooso et al. 2012; Chantaravisoot et al. 2015; Cai et al. 2020b). The exon corresponding to the hinge region of FLNB is frequently excluded in breast cancer and giant cell tumors (Li et al. 2018). Hinge exon exclusion is related to enhanced cell migration, the endothelial-to-mesenchymal transition and anti-apoptotic gene expression (van der Flier et al. 2002; Tsui et al. 2016; Li et al. 2018; Ma et al. 2020). RT-PCR results showed that the hinge exon of *FLNB* transcripts was frequently excluded in both low- and high- grade gliomas, again consistent with the Cuffdiff/Semi-Q analyses. The *FLNC* hinge exon was also excluded among the tested gliomas, albeit to a lesser extent than with *FLNB*, while the *FLNA* hinge exon was actually more frequently included in the high-grade gliomas. These patterns are also entirely consistent with the Semi-Q and Cuffdiff analyses, in which *FLNB* and *FLNC* hinge exon exclusion was detected in GBM, while *FLNA* was not found among the GBM-excluded CCEs. The basis for this differential frequency of hinge exon skipping between *FLNB, FLNC* and *FL*NA in gliomagenesis is unknown, but its significance is examined in more detail below.

GSN (gelsolin) and VILL (villin) are members of the Gelsolin/Villin superfamily. The two proteins are actin-modulating proteins that sever F-actin and cap the plus end of actin filaments (Li et al. 2012). It is also known that the two proteins can be cleaved by caspase-3 upon activation of apoptosis. The cleaved N-terminal fragment can function as a pro-apoptotic factor, while the C-terminal fragment functions as an anti-apoptotic factor (Kothakota et al. 1997; Azuma et al. 2000; Koya et al. 2000; Kusano et al. 2000; Sakurai and Utsumi 2006; Wang et al. 2016). Notably, exclusion of *GSN* exon 3, or exon 6 of *VILL*, generates N-terminal fragments with loss of pro-apoptotic function after cleavage, but C-terminal fragments remain intact and retain anti-apoptotic activity (Mazumdar et al. 2012). RT-PCR again confirmed the RNA-seq results, i.e., *GSN* exon 3 and VILL exon 6 were included more frequently in the normal samples, but largely excluded in the GBM samples, indicative of decreased apoptotic function.

CORO7 is an F-actin regulator involved in anterograde Golgi to endosome transport (Chan et al. 2011). Recently, it was shown that CORO7 functions as a positive regulator of the Hippo signaling pathway by interacting with a number of Hippo signaling proteins (Park et al. 2021). Hippo signaling can be activated by serum deprivation or high cell density, and activated Hippo downregulates proliferation and upregulates apoptosis (Masliantsev et al. 2021). Full-length CORO7 binds to several Hippo signaling proteins with high affinity, but truncated derivatives bind them with low affinity. Therefore, exclusion of exons 4 and 5 in GBM will likely downregulate Hippo signaling, thereby contributing to enhanced proliferation and reduced apoptosis. RT-PCR confirmed that these two exons were indeed predominantly skipped in GBM samples, again consistent with results from our RNA-seq analysis.

We also investigated whether the AS changes we detected in GBM, especially regarding the cell migration-related genes, were more or less represented in any of the three GBM subtypes, classical (CL), mesenchymal (MES), and proneural (PN). To this end, we first compared NHA samples with samples in each of the three GBM subtypes using Semi-Q, identifying CCEs as described above. The CCEs from each comparison with NHA were then compared with each other, and a large majority of events in each subtype, 70-80%, entirely overlapped (Fig. 2B). We then compared dysregulated CCEs in each of the three subtypes with CCEs from the comparison between NHA and all GBM samples (Fig. 2C, and Supplemental Table S6). Again, overlap was very high, ∼80-90%, with that between NHA and MES samples the highest. Interestingly, all of the 40 ‘actin cytoskeleton organization’ targets were included in the MES and PN subgroups, and all but four, *ACTN2, ACTN4, AGAP2* and *LBD3*, were included in the CL subgroup. These results indicate that differential AS in GBM is, with some exceptions, very similar in all three subtypes.

We also note that among the 107 GBM samples, we found two isocitrate dehydrogenase (IDH) mutant samples and three with unknown IDH status. Because IDH mutant glioma is not considered to be high-grade glioma (Najafi et al. 2022), we compared the NB and NHA samples with the remaining 102 GBM samples, using the Semi-Q method. As expected, the results were almost identical to those in our previous analysis: 96% of CCEs overlapped in the comparison between NB and GBM, and 99% of CCEs overlapped between NHA and GBM. Notably, all of the 40 cell migration-related genes that we have concentrated on were detected in this reanalysis.

### Differential gene expression analysis supports the appropriateness of comparing NHA and GBM samples

While we are concentrating here on GBM-specific differences in AS, we also examined gene expression, i.e., transcript abundance, in the NB, NHA and GBM samples. The ten NHA, 47 NB and 107 GBM samples analyzed above were therefore reanalyzed using DESeq2 (Love et al. 2014). Consistent with our findings above, principal component analysis (PCA) revealed that the NHA samples were clearly separated from the NB samples (Fig. 3A, upper left), and, significantly, that the NHA samples were closer to the GBM samples than were the NB samples (Fig. 3A, lower and right). These observations support our use of NHAs, rather than NB, to detect changes in AS in GBM.

**Figure 3.**
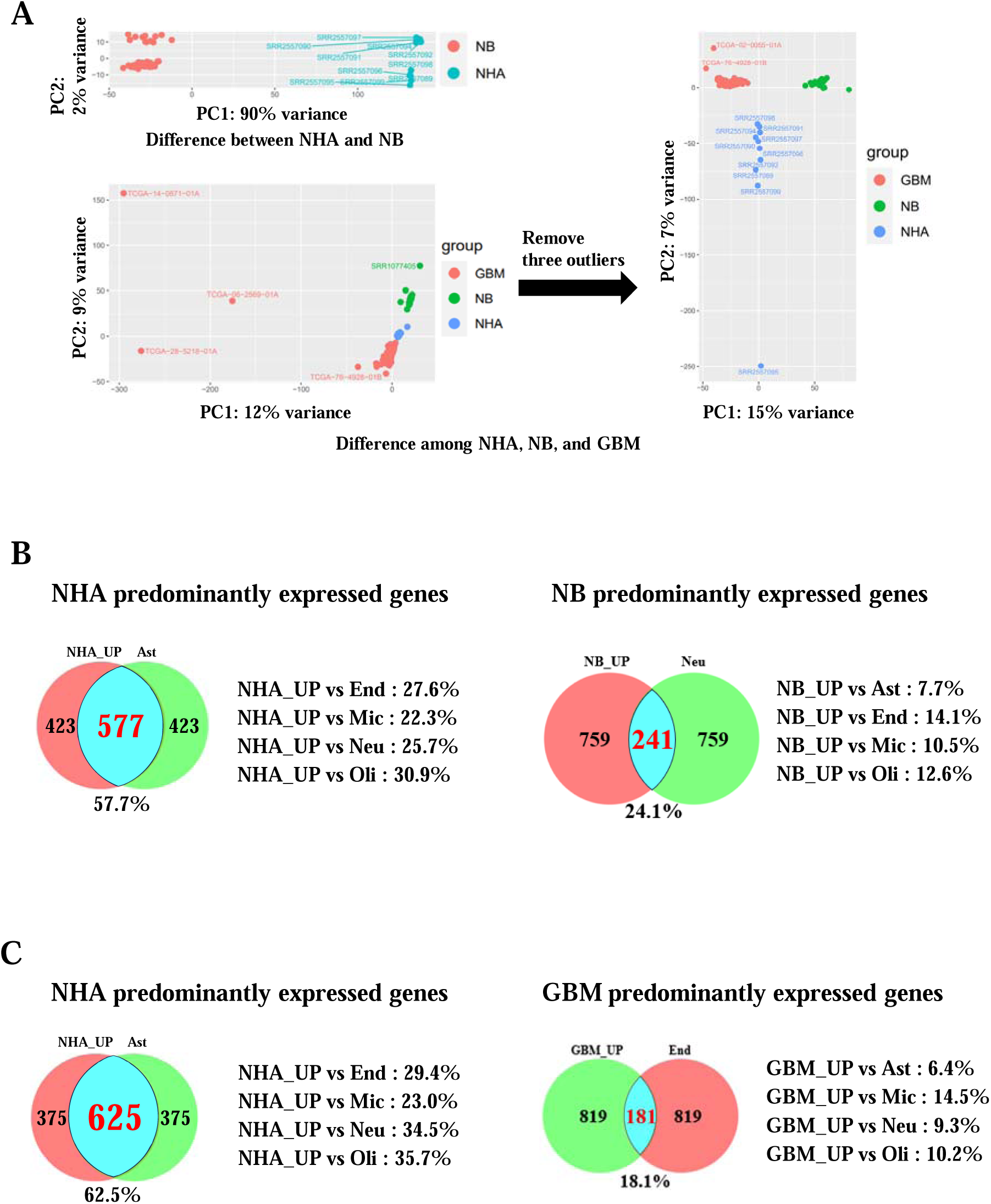
DGE analysis indicates that NHA samples are distinct from NB samples, and NB samples are closer to neurons than to astrocytes. A. Count files of NHA, NB and GBM samples were generated from BAM files by using htseq-count, and DGE analysis was performed using DESeq2. Comparisons between NB and NHA (upper left), between NB, NHA and GBM (lower left), and between NB, NHA and GBM with three outliers removed (right) are shown. The horizontal axis represents PC1 (Principal components 1, indicating the most variation in the data) and the vertical axis represents PC2 (Principal components 2, the second most variation in the data). **B**. 1000 genes that were upregulated in NHA and another 1000 genes that were upregulated in NB were identified by comparing NB with NHA. Each 1000 gene set was compared with previously reported brain subtype-specific upregulated genes. The number of genes and percent overlap are indicated. NB_UP most highly overlapped with neural-specific genes, and NHA_UP with astrocyte-specific genes (blue area and red lettering). Neu: neuron, Ast: astrocyte, Oli: oligodendrocyte, Mic: microglia, End: endothelial cells. **C**. 1000 genes that were upregulated in NHA (NHA_UP) and another 1000 genes that were upregulated in GBM (GBM_UP) were identified by comparing NHA with GBM. Each 1000 gene set was compared with previously reported brain subtype-specific upregulated genes. NHA_UP most highly overlapped with astrocyte specific genes, and GBM_UP with endothelial specific genes (blue area and red lettering). Neu: neuron, Ast: astrocyte, Oli: oligodendrocyte, Mic: microglia, End: endothelial cell.

We next wished to examine the cell-type specificity of the gene expression patterns in our samples. To this end, the top 1000 genes predominantly expressed in NB, NHAs and GBM, identified as described by McKenzie et al. (2018) using the above DESeq2 results (Fig. 3A), were compared with brain subtype-specific genes (Zhang et al. 2014; Zhang et al. 2016; McKenzie et al. 2018). Comparing NHA with NB, the genes predominantly expressed in NHAs as expected overlapped to the greatest extent with astrocyte-specific genes, while those expressed predominantly in NB overlapped to a high degree with neuron-specific genes (Fig. 3B). Comparing NHA with GBM, while the genes predominantly expressed in NHA again overlapped most frequently with astrocyte-specific genes, genes expressed predominantly in GBM displayed the highest overlap with endothelial-specific genes (Fig. 3C). These results are consistent with observations that GBM stem cells often lose their original glial characteristics and transdifferentiate into vascular endothelial-like cells, which can enhance angiogenesis in the tumor mass (Ricci-Vitiani et al. 2010; Wang et al. 2010; Soda et al. 2011; see below for further discussion). Notably, when NB and GBM were compared, neuron-specific genes were again the most prominently expressed in NB, but were very lowly expressed in GBM (Supplemental Fig. S9). These findings further support our conclusion that the apparent GBM-specific microexon exclusion we observed in our initial comparison of NB with GBM reflected the presence of neurons in the NB but not GBM samples, and thus was not due to a change in AS that occurs during gliomagenesis. These findings further highlight the validity of comparing splicing patterns in NHAs and GBM as a measure of AS dysregulation in GBM.

### Filamin hinge-exon exclusion increases expression of genes relevant to gliomagenesis

As shown above, *FLNB* and *FLNC* (although not *FLNA*) undergo increased hinge exon exclusion in GBM, with *FLNB* showing the largest change relative to NHA cells. This should result in less filamin cleavage, less translocation of cleaved fragments to the nucleus, and hence altered transcriptional activity (Bedolla et al. 2009; Deng et al. 2012; Mooso et al. 2012; Savoy and Ghosh 2013). To investigate this possibility, we utilized previously described *FLNB* hinge exon-deleted (ΔH) human mammary epithelial cells (HMLE) and control, unedited HMLE cells (Li et al. 2018), isolated RNA from them and performed RNA-seq (Fig. 4A, PCR confirming hinge-exon deletion; 4B, PCA displaying variance between samples; and Supplemental Fig. S6, additional statistical analyses). We compared the two groups using DESeq2 and determined the top 50 upregulated and top 50 downregulated genes in the FLNBΔH group (Supplemental Table S7). GO analysis revealed no terms related to biological processes in the downregulated genes, but terms related to cell proliferation and anti-apoptotic processes characterized the upregulated genes (Fig. 4C and 4D). RT-qPCR confirmed that all 18 genes in these two categories were indeed upregulated in the FLNB-HMLE cells (Fig. 4E and Supplemental Table S8; note that one gene, *SOD2*, was discarded because of technical difficulties with PCR amplification). Notably, our DESeq2 analysis described above found that 15 of these 18 genes were also upregulated in GBM.

**Figure 4.**
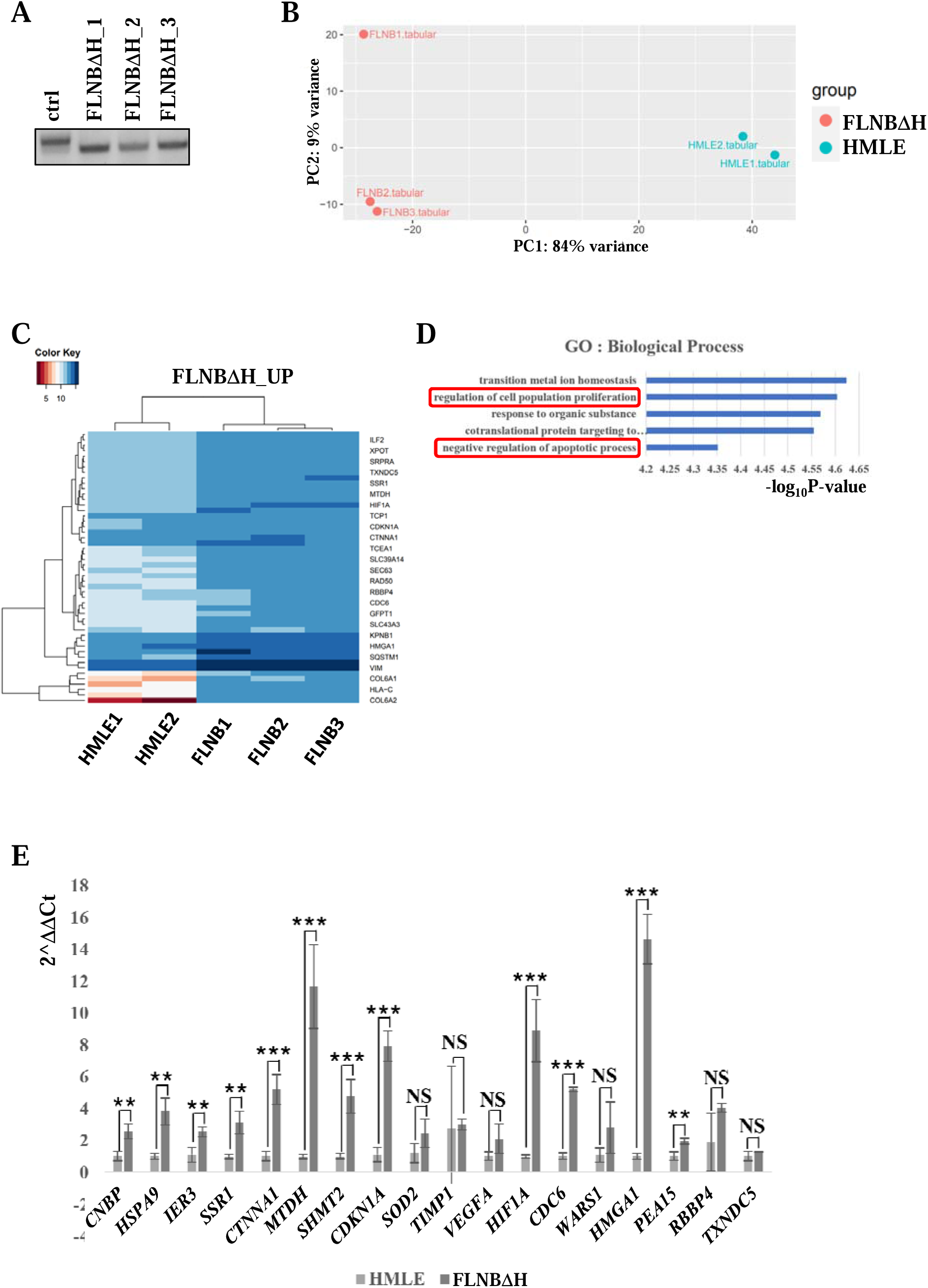
Hinge exon deletion in *FLNB* dysregulates expression of genes involved in cell growth. A. *FLNB* hinge exon-deleted HMLE cells and control Cas9-HMLE cells were analyzed by PCR to confirm hinge-exon deletion. Hinge-nested exon specific primers as described in Figure 2A were used, and one control and three different hinge exon-skipped clones were analyzed. B. Counts files were generated from fastq files using the kallisto-quant program, and DGE analysis was performed using DESeq2. The PCA plot is drawn in the same manner as in Figure 3A. C. The top 50 upregulated genes in FLNBΔH compared to control cells were selected and are shown in the heatmap. The horizontal axis indicates samples analyzed and the vertical axis displays the 50 most upregulated genes (which are all listed in Supplemental Table S7). D. GO analysis using the 50 most upregulated genes was performed and the top categories are indicated. Terms related to cell migration, proliferation and apoptosis are highlighted with rectangles. The horizontal axis represents minus log10 of p value. E. Upregulated genes from the groups “regulation of cell population proliferation” and “negative regulation of apoptotic process” were analyzed by RT-qPCR using RNA isolated from *FLNB* hinge-deleted HMLE cells and control HMLE cells. The vertical axis indicates 2^ΔΔCt value and the horizontal axis the genes analyzed. * p<0.05, ** p<0.01, *** p<0.001 and NS, not significant. Error bars represent the standard deviation of each data set. n=4 repeats.

To extend these analyses to *FLNA* and *FLNC*, we used CRISPR/Cas9 to create hinge exon-deleted HeLa cell lines and controls, and again carried out RNA-seq analysis (Fig. 5A, B, and Supplemental Fig. S7 for FLNC and Fig. 5E, F, and Supplemental Fig. S8 for FLNA; note that since the experiments with FLNB and FLNA/C utilized different cell lines, they may not be directly comparable). We again compared the two groups using DESeq2 and determined the top 50 upregulated and top 50 downregulated genes in both the FLNAΔH and FLNCΔH groups (Supplemental Table S7). GO analysis revealed no terms related to biological processes in the upregulated genes, but there were a number of terms related to mitochondrial function that characterized the downregulated genes (Fig. 5C and Supplemental Fig. S4A for FLNC, and Fig. 5G and Supplemental Fig. S4C for FLNA). For FLNCΔH, RT-qPCR confirmed that *ND6, TCERG1* and *TRMU* were downregulated (Fig. 5D, Supplemental Fig. S4B and Supplemental Table S8), while for FLNAΔH, *AURKB, ATP6, CO1, CYB, ND1, ND4* and *TRMU* were confirmed by RT-qPCR to be downregulated (Fig. 5H, Supplemental Fig. S3D and Supplemental Table S8). It is notable that two of the three FLNCΔH- downregulated genes and five of the seven FLNAΔH-downregulated genes were also dysregulated in our NHA vs GBM comparison, although the significance of this with FLNA is unclear since FLNA hinge exon skipping in GBM was minimal (see above).

**Figure 5.**
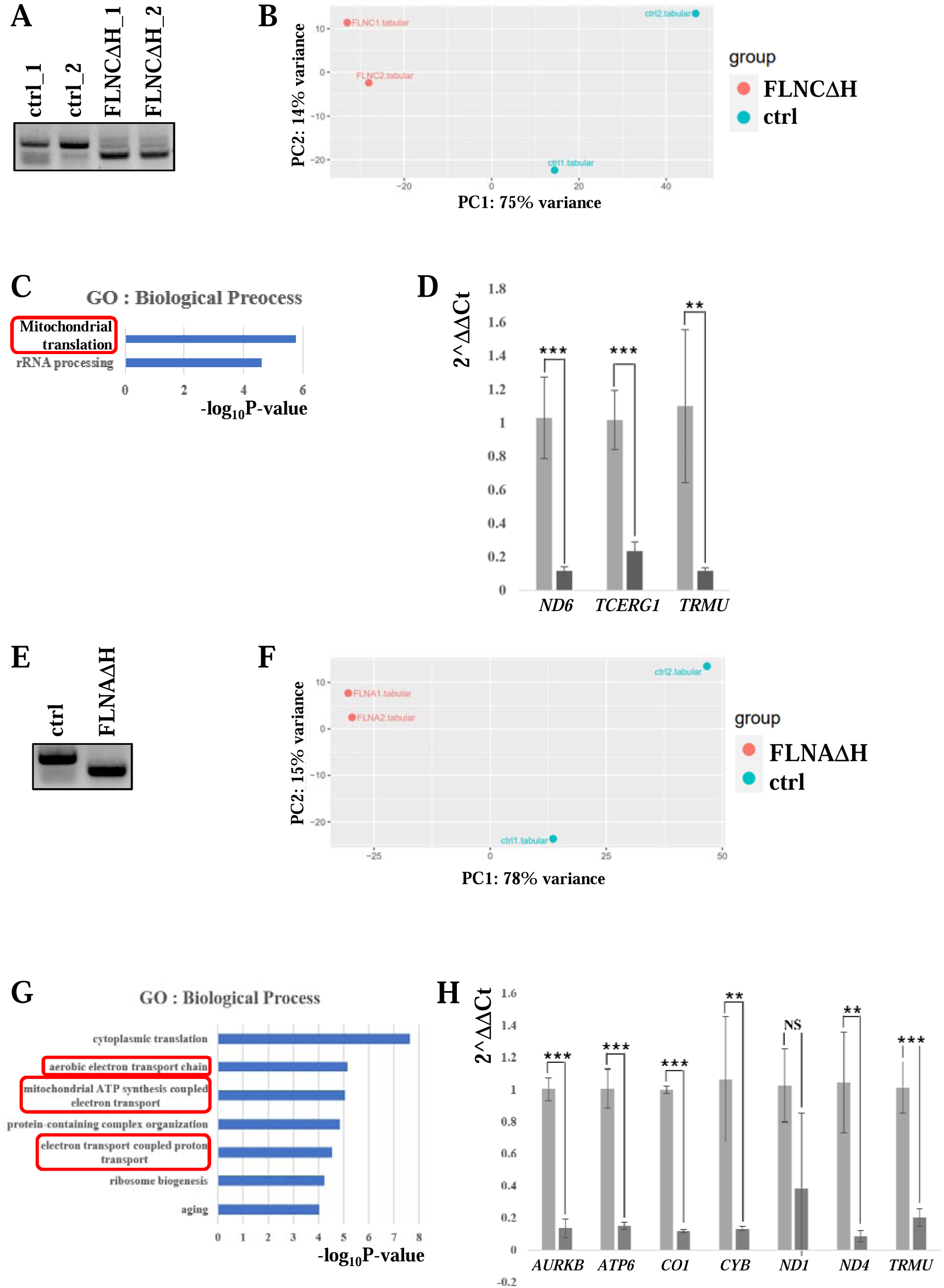
Hinge exon deletion of *FLNA* and *FLNC* also dysregulates gene expression. **A**. Hinge exon-deleted *FLNC* HeLa cells (FLNCΔH) and control HeLa-Cas9 cells were generated by CRISPR as described in materials and methods. PCR validation of two independent control and two independent FLNCΔH clones was performed by using hinge-nested exon specific primers as described in Figure 2A. **B**. Counts files were generated from fastq files using the kallisto-quant program, and DGE analysis was performed using DESeq2. The PCA plot is drawn in the same manner as in Figure 3A. **C.** The top 50 downregulated genes in FLNCΔH were selected, and GO analysis was performed. The results revealed that *FLNC* hinge exon deletion is significantly related with dysregulation of mitochondrial translation (red rectangle). The horizontal axis represents minus log10 of p value. **D**. RT-qPCR was performed with RNA from control and hinge-deleted FLNC HeLa cells for the three genes, *ND6, TCERG1* and *TRMU*, in the group ‘translation’ (red rectangle in Fig. 5C). * p<0.05, ** p<0.01, *** p<0.001. Error bars represent the standard deviation of each data set. n=4 repeats. **E**. Hinge-deleted *FLNA* HeLa cells (FLNAΔH) and control HeLa-Cas9 cells were generated by CRISPR as described in materials and methods. PCR validation was performed by using hinge-nested exon specific primers as described in Figure 2A. **F.** Counts files were generated from fastq files by using the kallisto-quant program, and DGE analysis was performed using DESeq2. The PCA plot is drawn in the same manner as in Figure 3A. **G.** The top 50 downregulated genes in FLNAΔH were selected, and GO analysis was performed. The results revealed that hinge exon deletion of *FLNA* is significantly related with dysregulation of mitochondrial function (red rectangles). The horizontal axis represents minus log10 of p value. **H**. RT-qPCR was performed with RNA from control and hinge-deleted *FLNA* HeLa cells for the seven genes, *AURKB, MT-ATP6, CO1, CYB, ND1, ND4* and *TRMU*, in the group related to mitochondrial function (red rectangles in Fig. 5G). * p<0.05, ** p<0.01, *** p<0.001 and NS, not significant. Error bars represent the standard deviation of each data set. n=4 repeats.

### Hinge exon exclusion of FLN A, B and C alters cell growth properties

We next examined the FLNBΔH HMLE cells to determine whether they behaved in culture in a manner consistent with suggestions from previous studies and with the changes in gene expression described above. First, we examined the morphology of the control and FLNBΔH HMLE cells. Strikingly, morphological features of the two cell lines were quite different (Fig. 6A). After 24 hours of subculturing, the control cells were not tightly attached to the bottom of the cell culture dish and exhibited round-shaped morphology, while the FLNBΔH cells attached tightly to the dish and exhibited fibrous morphology, which is related to a high rate of cell migration (Nagano et al. 2012; Tojkander et al. 2012). To measure cell migration rates more directly, we performed a cell infiltration assay (see Materials and methods). The FLNBΔH cells exhibited twice the migration rate as control cells when serum was present in the bottom chamber, and three times the rate when an inducer of cell migration, supplied by the manufacturer, was present in the bottom chamber (Fig. 6B and Supplemental Table S9). Next, we performed MTT assays to compare proliferation of the two cell lines. The results revealed that the FLNBΔH cells exhibited a 1.5-2.0 fold higher proliferation rate compared with control cells after five days of culture (Fig. 6C and Supplemental Table S10). Finally, we examined sensitivity to ionizing radiation as a measurement of the apoptotic cell death propensity of the two cell lines (Fig. 6D and Supplemental Table S11). 72 hours after exposure to 20Gy of ionizing radiation, flow cytometry (upper panels, quantitation on lower panel) revealed that annexin-positive HMLE-Cas9 control cells increased by 2.0 fold, but annexin-positive FLNBΔH cells increased by only 1.4 fold, a small but significant decrease (p=0.0077). It is also notable that even in the absence of irradiation, the percentage of annexin-positive FLNBΔH HMLE cells was markedly less than observed with the control HMLE cells (3.5% vs 13.6%), which is in agreement with previous findings (Ma et al. 2020). Together, these results are consistent with the changes in gene expression we observed in FLNBΔH HMLE cells, and also with known properties of GBM cells (Lefranc et al. 2005; Delgado-Martín and Medina 2020).

**Figure 6.**
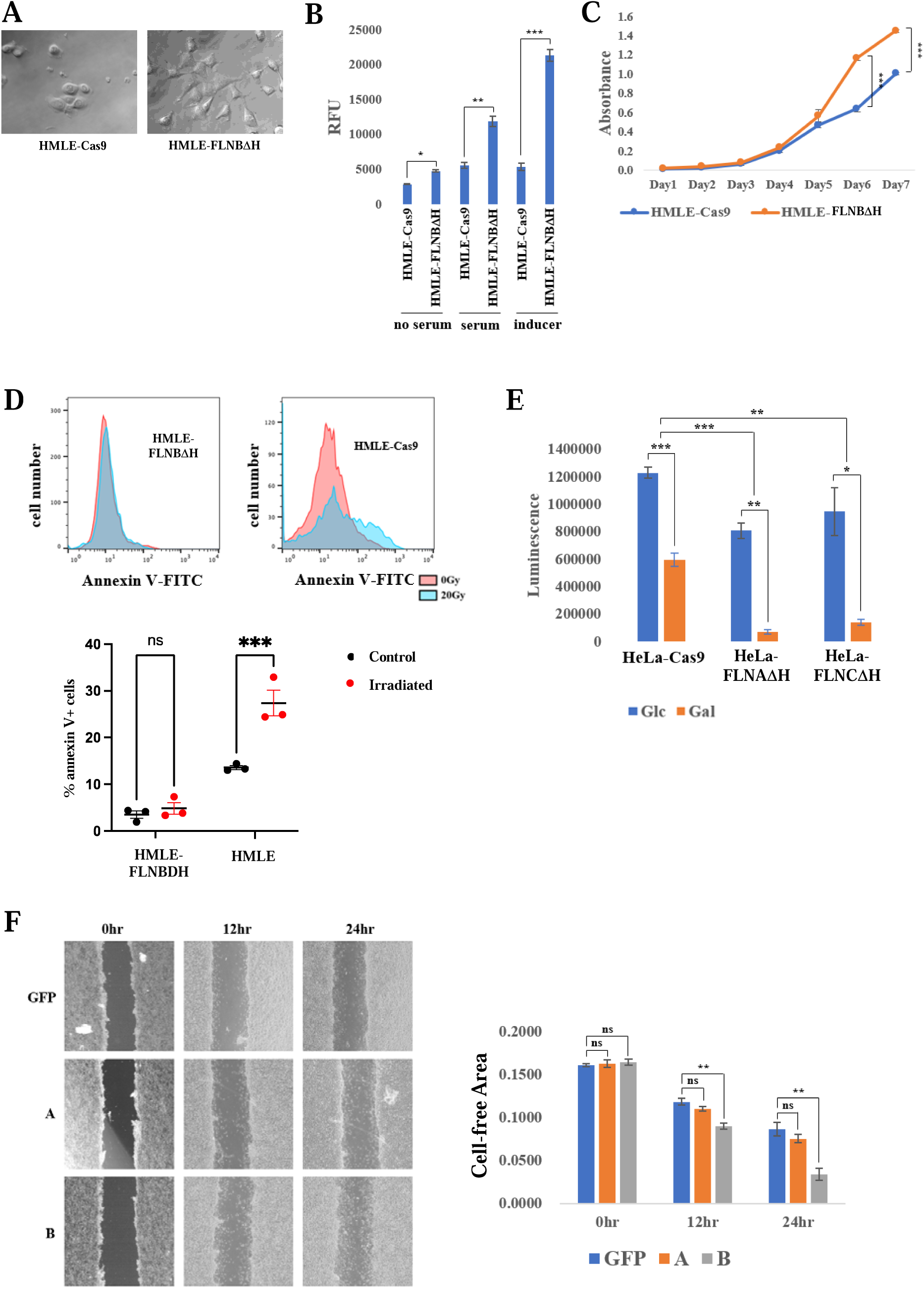
GBM-specific AS of FLN/ACTN transcripts alters cellular growth properties. **A**. FLNBΔH and control HMLE-Cas9 cells display different morphologies. Cells were subcultured at a density of 1X10^6^ per 100mm dishes and cells were imaged and photographs taken 24 hours later at a magnification of 100X. **B**. FLNBΔH cells migrate faster than control HMLE-Cas9 cells. Transwell infiltration analysis was performed as described in materials and methods, in the presence of no serum, serum or inducer as indicated. Migrated cells were stained with fluorescent dye and the relative fluorescence of samples was measured. The vertical axis represents relative fluorescence units (RFU). * p<0.05, ** p<0.01, *** p<0.001. **C.** FLNBΔH cells are more proliferative than control HMLE-Cas9 cells. MTT assays were performed as described in materials and methods. Cells were cultured at a density of 5000 per well in 24 well plates and grown for the number of days indicated. Total proliferation was measured by absorbance at 560nm (vertical axis). **D**. FLNBΔH cells are resistant to apoptosis. Control HMLE-Cas9 and FLNBΔH cells were cultured and irradiated at 0 Gy or 20 Gy as indicated. After irradiation, cells were cultured for an additional 72 hours, and annexin V staining and flow cytometry analysis was performed as described in the materials and methods. Vertical axis represents cell number and horizontal axis represents intensity of the Annexin V-FITC signal (upper panel). Percent annexin V-positive cells are indicated on the vertical axis (lower panel). **E**. Mitochondrial catabolic function is dysregulated in both FLNAΔH and FLNCΔH cells. Mitochondrial catabolic function was measured as described in materials and methods. The ATP production difference was compared between two growth conditions, cells cultured in glucose-containing (Glc) or galactose-containing (Gal) media. Total ATP production was measured by luminescence (vertical axis). **F.** Exon 8B inclusion in *ACTN4* enhances cell migration. pEGFP-C1 (GFP), GFP-tagged ACTN4A (A) and GFP-tagged ACTN4B (B) encoding plasmids were transfected into A172 cells. Wound healing analysis was performed as described in the materials and methods. After 0, 24 and 48 hours, images of wound area were captured by microscopy at 40x magnification (left). Wound areas were calculated using ImageJ (right).

Hinge exon skipping in FLNA and FLNC transcripts brought about changes in gene expression related to mitochondrial catabolic functions. To investigate the possible significance of these findings, which would reflect changes known to occur in GBM (Lefranc et al. 2005; Lefranc et al. 2018; Delgado-Martín and Medina 2020), we performed mitochondrial ATP-production assays using the control, FLNAΔH and FLNCΔH HeLa cells described above. While ATP production in the control cells was decreased ∼3 fold when glucose-containing media was changed to galactose-containing media, it was reduced ∼8 fold and ∼5 fold in the FLNAΔH and FLNCΔH cells, respectively, following the change (Fig. 6E and Supplemental Table S12). As a control, we also analyzed mitochondrial anabolic function, using an MTT assay. The results (Supplemental Fig. S5 and Supplemental Table S13) revealed no differences among the control, FLNAΔH and FLNCΔH cells, supporting the specificity of our results.

### GBM-related ACTN4 AS enhances cell migration

As described above, our data revealed that *ACTN4* (and *ACTN2*) AS is dysregulated in GBM, resulting in increased inclusion of exon 8B relative to 8A. As mentioned, actinins function to enhance stress fiber formation and cell migration, and enhanced exon 8B inclusion has been shown to increase stress fiber formation in lung cancer (Honda et al. 2004). To extend these findings to GBM, we measured the effects of ACTN4 splice isoforms on cell migration in A172 GBM cells. To do so, GFP-tagged ACTN4A (containing exon 8A) and ACTN4B (containing exon 8B) were cloned into pFLAG-CMV7.1, and, together with a GFP-only control, expressed in A172 cells. We then performed wound-healing assays after the cells reached confluency. The total wound cell-free area was estimated, and relative ratio changes were calculated. While the GFP-ACTN4A-expressing cells exhibited almost the same ratio change compared to the GFP-only control group, the GFP-ACTN4B-expressing cells exhibited more than twice the change compared to the controls (Fig. 6F and Supplemental Table S14). As with our findings regarding filamins, these results strengthen the view that dysregulated AS in GBM contributes to tumorigenesis.

## Discussion

In this paper, we developed and validated Semi-Q, a new, simplified method for AS analysis. Using Semi-Q, we found that NHAs offer advantages over NB for comparative AS analysis with GBM. Comparison of NHA with GBM revealed that cell migration-related genes are among the most frequently dysregulated by AS in GBM, and that the GBM gene expression pattern is closer to that of endothelial cells than to astrocytes. We found that splicing of transcripts encoding filamins and actinins, among others, is substantially altered in GBM, and that these changes produce protein isoforms with properties that are closely related to gliomagenesis, including enhanced cell migration and decreased cell death. Below we discuss these findings both with regards to our computational pipeline and from the perspective of how AS contributes to GBM.

To facilitate analysis of AS in hundreds of samples, we developed the method we refer to as Semi-Q. As part of the validation process, we compared results obtained using Semi-Q with the established but more time-consuming methods Cufflinks-Cuffdiff and rMATS (Liu and Rabadan 2021). Interestingly, we found that Semi-Q, and in some respects Cufflinks-Cuffdiff (CC), are more useful and reliable than rMATS in analyzing AS changes, from three perspectives. First, in the comparison between NHA and GBM, inclusion of microexons was far more pronounced with Semi-Q and CC than with rMATS. Semi-Q and CC both revealed that ∼10% of the total NHA-specific CCEs are microexons, which decreased slightly in GBM, while rMATS found that <0.5% of CCEs are microexons, in both NHA and GBM. The results of CC and Semi-Q well-represent known levels of microexon inclusion in glial cells and exclusion in GBM, but rMATS does not. Moderate levels of microexon inclusion in astrocytes is well-established, and exclusion of glial microexons, detected using event-based programs, is thought to reflect a dedifferentiation process of glioma cells (Irimia et al. 2014; Head et al. 2021; Reixachs-Solé et al. 2020).

Second, comparing the nature of the CCE-containing transcripts, we observed that expression signatures of neuronal cells in the NB samples and immune cells in the GBM samples were more pronounced in the Semi-Q results than with either of the other two methods. CCEs are tissue-specifically included, and patterns of inclusion/exclusion are characteristic of different cell types (Clark et al. 2007; Ellis et al. 2012). For example, microexon inclusion is to a significant degree neuron-specific, and included microexons can change the function of proteins towards neural-specific functions (Irimia et al. 2014). Thus, our detection of microexons and other neural-specific CCEs in the NB samples with Semi-Q is highly indicative of neurons, which is of course as expected in whole-brain samples (Irimia et al. 2014; Ustianenko et al. 2017; Gonatopoulos-Pournatzis and Blencowe 2020; Lee et al. 2022). Similarly, our detection with Semi-Q of immune-function related CCEs in the GBM samples is indicative of the presence of immune cells. As mentioned above, there is considerable evidence for the presence of such cells in GBM tumors, and infiltrating immune cells enhance inflammatory responses and are related to poor prognosis (Komohara et al. 2008; Quail et al. 2016; Quail and Joyce 2017; Zhang et al. 2017). Our comparison by Semi-Q of CCE-containing transcripts between two groups coupled with GO analysis was thus able to reveal previously observed heterogeneity of tissue samples, as we have done here with NB and GBM.

Third, the CCE-containing genes that were validated by RT-PCR were included in the Semi-Q and CC results, but not in the rMATS results. Enhanced cell migration and proliferation, resistance to cell death and downregulation of mitochondrial function are features of GBM (Lefranc et al. 2005; Lefranc et al. 2018; Delgado-Martín and Medina 2020). Semi-Q was thus indeed able to identify CCE-containing transcripts that are subject to disease-relevant AS in GBM. Together, our results indicate that Semi-Q will be a valuable tool in deciphering tissue- and disease- specific AS patterns.

Our analysis, using Semi-Q and CC, identified over 500 genes that produce transcripts that display altered splicing in GBM. We investigated here genes identified by GO analysis as being related to cell migration, and specifically encoding proteins involved in actin cytoskeleton organization. GBM cells are highly infiltrative, characterized by aggressive tumor cell migration and invasion (Lefranc et al. 2018). While the factors contributing to this are complex, changes in the actin cytoskeleton play a significant role (Nagano et al. 2012; Masoumi et al. 2016; Zottel et al. 2021). Indeed, the genes/gene families we identified and studied here have all been implicated in cancer, and the GBM-enhanced AS events we uncovered all result in protein isoforms with functions that would increase tumorigenicity, based on previous studies and/or our own results. Notably, several earlier studies linked changes in expression of cell migration-related genes, including those encoding actinins, filamins and gelsolins, specifically with GBM (Rickman et al. 2001; Sen et al. 2009; Chantaravisoot et al. 2015; Kamil et al. 2019), while another linked a high-grade pediatric glioma with an altered filamin arising from a splice site mutation in *FLNA* (Marinari et al. 2021). Additionally, a member of the GSN/VILL family, AVIL, was found to be overexpressed and a driver of tumorigenesis in GBM (Xie et al. 2020; Cornelison et al. 2021).

The mechanisms underlying the splicing changes we observed remain to be determined, but likely involve alterations in expression of splicing regulators. While splicing dysregulation in cancer frequently involves mutations in splicing factors (Bradley and Anczuków 2023), this seems unlikely to be the case in GBM (Braun et al. 2017). Indeed, several studies identified hnRNPs, notably hnRNP A1/A2 and PTBP1, as being upregulated in GBM, altering splicing of specific transcripts that contribute to gliomagenesis in various ways, including enhanced cell proliferation and migration (Cheung et al. 2009; David et al. 2010; Golan-Gerstl et al. 2011; Deng et al. 2016). Overexpression of these proteins is driven transcriptionally by elevated levels of the oncogenic transcription factor c-Myc (David et al. 2010) and in the case of PTB1 also by gene amplification and downregulation of a PTBP1-targeting miRNA (Ferrarese et al. 2014). However, a large-scale RNA-seq analysis revealed that expression of at least 58 RNA binding proteins (RBPs), including for example several SR proteins, is altered in GBM and GSCs (Correa et al. 2016). Indeed, various RBPs have been shown to contribute to stemness, migration and invasion of glioma cells (Velasco et al. 2019). Among them, deficiency of the RBP Quaking (QKI) has been implicated in maintenance of GSC stemness and thus contributes to GBM invasiveness (Shingu et al. 2017; Han et al. 2019). Notably, QKI functions in AS regulation of both *ACTN2* exon 8A/B (Chen et al. 2021; Montañés-Agudo et al. 2023) and the *FLNB* hinge exon (Li et al. 2018).

Several previous studies have examined global AS changes in GBM. These have used multiple different methods, usually junction-based, and have typically compared GBM with NB (Correa et al. 2016; Babenko et al. 2017; Li et al. 2019; Song et al. 2019; Zhou et al. 2019; Wang et al. 2021). However, RNA-seq and GO analyses in these studies did not reveal significant differential AS of cell migration-related genes between NB and GBM. This could reflect the methods employed and/or the use of NB for comparison with GBM. In fact, our analyses using NB for comparison also failed to reveal cell migration-related genes as a prominent category for misspliced transcripts; they likely were obscured by the large presence of neuronal cells/transcripts in these samples. Several other studies, however, found that AS of specific transcripts, or the presence of novel isoforms, can promote cell migration in GBM, but analyses were limited to individual genes (Yu et al. 2007; Lo et al. 2009; Golan-Gerstl et al. 2011; LeFave et al. 2011; Uceda-Castro et al. 2022). Interestingly, one such study, using the program AS detector, observed that certain cell migration-related transcripts are misspliced in GBM (Zhou et al. 2019). Among them, MYO1B, CTTN and KTN1 overlapped with our results, and the dysregulated exons detected were identical. However, to our knowledge, ours is the first study to identify misspliced actin/cytoskeletal-related transcripts, such as actinins and filamins, and to show that the encoded protein isoforms all favor enhanced cell migration and proliferation. While for technical reasons our experiments examining the functions of the FLN hinge exon deleted isoforms could not be performed in GBM-derived cells (see Materials and methods), the effects we observed all reflect properties known to be important for gliomagenesis (e.g., Cai et al. 2020b, Chantaravisoot et al. 2015, Kamil et al. 2019).

Our differential gene expression analysis also provided insights into significant changes that occur during gliomagenesis. Glioblastoma stem cells (GSCs) can transdifferentiate into endothelial cells, and GSC-derived endothelial cells (ECs) are important regulators of angiogenesis and vasculogenesis (El Hallani et al. 2010; Ricci-Vitiani et al. 2010; Soda et al. 2011; Scully et al. 2012; Mei et al. 2017). GSC-derived ECs proliferate excessively, and construct loosely networked excessive vessel branching. Tumors not only acquire blood supply through these vessels, but also penetrate into loose EC networks and migrate to new locations. GSCs also differentiate into EC-like cells, via the process of vasculogenic mimicry (VM; Maniotis et al. 1999). VM structures are not functional endothelial vessels but hollow channel structures in which functional microcirculation is observed. CD31 and CD34, which are only expressed in ECs and not VMs (Maniotis et al. 1999), are the most prominent markers of differentiating EC-derived vessels (Maddison et al. 2023). Notably, in our comparison between NHA and GBM, *CD31* and *CD34* were not included among the genes predominantly expressed in our GBM samples, although a number of genes encoding known EC (and VM)-markers, including *CD44, ENG, ACTA2, MMP2, VWF* and *TNFRSF21* (Maddison et al. 2023), were prominent among the GBM-enriched genes. Our DGE analysis thus indicates that the GBM samples we analyzed displayed VM-associated characteristics. VM is associated with high-grade gliomas, as well as with tumor progression, invasion, infiltration and poor prognosis (Ricci-Vitiani et al. 2010; Soda et al. 2011; Cai et al. 2020a), which are all properties that characterize GBM.

In summary, we developed a novel computational pipeline, Semi-Q, and used it to analyze changes in alternative splicing in GBM. Our results provide novel insights into the high motility and invasiveness of GBM and thereby both shed new light on our understanding of the disease, and also provide considerable evidence that altered AS contributes to disease progression.

## Materials and Methods

### Selection of samples

90 NB samples were downloaded from the Genotype-Tissue Expression (GTEx) Project in the dbGaP database, and 155 GBM samples were downloaded from The Cancer Genome Atlas (TCGA) in the Genomic Data Commons database (Supplemental Table S1). Eleven NHA RNA-seq datasets were downloaded from the Sequence Read Archive (SRA) database (Supplemental Table S1). One NB sample was deleted because it did not align with the human genome.

To ensure an accurate comparison between these samples, we first selected RNA-seq data of high quality by Gene Body analysis, in which the mapped reads displayed minimal bias towards the 3’- vs 5’- end along the gene transcripts (i.e., with minimal degradation in the gene body) (Wang et al. 2012). We set a coverage ratio of 65% (ratio of reads mapped to the 5’-end half vs those to the 3’-end half of gene transcripts) as cutoff and identified 47 NB and 143 GBM high quality samples (Supplemental Fig. S1 and Supplemental Table S2).

To focus on high-grade glioma samples, we developed the following protocol. Among the selected 143 GBM samples, three were unclassified, eight were glioma CpG island methylator phenotype (G-CIMP), 35 were classical, 47 were mesenchymal, 26 were proneural and 24 were neural subtypes. The three unclassified samples were discarded, because we were interested in clearly classified high-grade glioma samples. The eight G-CIMP samples were discarded as well, because G-CIMP is a distinct subset of gliomas, and are more prevalent among lower-grade gliomas (Noushmehr et al. 2010). The 24 neural subtype samples were also discarded, because they could arise from contamination of neural tissue (Sturm et al. 2012; Gill et al. 2014; Behnan et al. 2017).

The 47 NB and remaining 108 GBM samples were analyzed further. First, they underwent an alignment process using the STAR mapping program (Dobin et al. 2013). After running STAR, one GBM sample was discarded because of low uniquely mapped reads percentages. For the NHA samples, one sample was discarded because of small size (370MB) and because the fastq quality check was poor. 47 NB samples, 107 GBM samples, and ten NHA samples were thus selected for further analysis.

### RNA-seq analysis

#### Identification of CCEs

NB and GBM data were downloaded in BAM file format, and NHA in SRA format. FASTQ files were generated from the downloaded BAM files and SRA files using bedtools and fastq-dump commands, respectively. All the FASTQ files were analyzed for quality using the FASTQC program, after which BAM files were generated using the STAR alignment program (Dobin et al. 2013). The gtf and fasta files were downloaded from the NCBI refseq database. The BAM files underwent sorting and indexing using the samtools program. For the Cufflinks-Cuffquant-Cuffdiff procedure, the sorted and indexed bam files were assembled using Cufflinks, merged with Cuffmerge, and quantified using Cuffquant and Cuffdiff (Trapnell et al. 2012). Finally, using the splicing.diff and cds_exp.diff files, isoforms were compared and CCEs were obtained. We collected genes whose significance value was ‘yes’, and the JSD value was higher than 0.2 in the splicing.diff file, and collected isoforms with status value ‘OK’ in the cds_exp.diff file.

For the Semi-Q procedure, FPKM values were generated and converted into percentage FPKM values per gene after the indexed bam files were assembled using Cufflinks. By using a p-value cutoff of 0.01 and a difference ratio value cutoff (ΔFPKMr) of 0.2, oppositely regulated genes between normal and GBM were identified (Supplemental Fig. S2A and S2B), and these isoforms were compared using the IGV program (Supplemental Fig. S2C). The Semi-Q procedure is explained in more detail in Supplemental Fig. 2.

#### DGE analysis

For NB, NHA and GBM samples, we used a gtf file from the Ensembl database and the htseq-count program, and for the analysis of hinge-exon skipping of filamins, we used the fasta file from the Ensembl database and the kallisto program. The count tables generated from the htseq-count and kallisto were analyzed using DESeq2, and the files were used to draw PCA plots and heatmaps (Love et al. 2014). We set a 0.05 p-value and 100 counts difference as cutoff, i.e., by collecting the top 50 genes whose difference value was higher than 100 (upregulated in ΔH) or the 50 less than -100 (downregulated in ΔH).

### CRISPR-editing FLNAΔH- and FLNCΔH-HeLa cells

For FLNA hinge-KO, the sequences 5’-TGGAGAGTCTGTTGTCACAG-3’ (front) and 5’- CTAGTTTACCATGTGTGAGG-3’ (rear) were selected as target sequences. For FLNC hinge-KO, the sequences 5’-GTAGCCACAGCAAGACTAGA-3’ (front) and 5’- AAGGAGTGCTGTGCCCAAGT-3’ (rear) were selected as target sequences. The oligomers were cloned into the PX459-V2.0 vector (Cong et al. 2013). HeLa cells were seeded at ∼50% density in 100mm culture dishes, and plasmids were transfected. Two days later, transfected cells were counted and transferred to 96-well plates at single cell density. All the cells were grown to confluence. Hinge KOs were confirmed by RT-PCR. For unknown reasons, we were unable to obtain FLN hinge exon KOs with U87 or A172 GBM cells.

### Cells, plasmids and transfection

GFP-tagged ACTN4A and ACTN4B-expressing plasmids were kindly donated by Prof. Yamada at the National Cancer Center Research Institute, Tokyo, Japan (Miyanaga et al., 2013). HeLa, CRISPR-edited HeLa, A172, U87 and NHA cells were cultured in DMEM supplemented with 10% FBS. HMLE and CRISPR-edited HMLE cells were kindly donated by Prof. Hahn, Harvard Medical School, Boston, United States (Li et al. 2018). The HMLE and CRISPR-edited HMLE cells were cultured in MEGM^TM^ mammary epithelial cell growth medium BulletKit^TM^ (Cat. No. CC-3150, Lonza). All cells were cultured at 377#x00B0;C in a 5% CO_2_ incubator. Transfections were all performed with a Neon Electroporation System (Cat. No. MPK5000S, Invitrogen) following the manufacturer’s instructions.

### RNA purification, RT-PCR and qPCR, and RNA-seq

All the NB, low-grade and high-grade glioma patient samples were obtained from the Herbert Irving Comprehensive Cancer Center Tumor Bank (Supplemental Table S1). For RT-PCR, samples were dissolved in 1ml of TRIzol^TM^ reagent (Cat. No. 15596018, Invitrogen), and purification was performed according to the manufacturer’s instructions. Purified RNA was dissolved in RNAse-free water, and 1µg of each RNA was used for reverse transcription with Maxima Reverse Transcriptase (Cat. No. EP7042, Thermo Fisher) using oligo(dT)-20mer. All cultured cells (NHA, A172, and U87) were treated and processed in the same way as tissue samples. For RT-PCR, 1µl of total cDNAs were used for template, and PCR reactions were performed using EconoTaq® PLUS GREEN 2X Master Mix (Cat. No. 30033-2, Biosearch^TM^ Technologies). Total PCR reaction samples were loaded on 1% agarose gels (Cat. No. BP160-500, Fisher Bioreagents) for analysis of standard CCEs. Products were quantified by ImageJ, and average values of two replicates are shown. For analysis of microexon inclusion, 3% MetaPhor^TM^ Agarose gels (Cat. No. 50181, Lonza) were used. Experiments were repeated multiple times (>3) with equivalent results to those shown. For real-time qPCR, 1:100 diluted cDNAs were used for template, and PCR reactions were performed using the Power SYBR® Green PCR Master Mix (Cat. No. 4367659, Applied Biosystems) with the StepOnePlus qPCR system (Applied Biosystems). qPCR data were analyzed using the -ΔΔC_T_ method. Averages from four independent replicates are shown. Sequence information for primers is in Supplemental Table S15.

For RNA-seq analysis of CRISPR-edited HeLa and HMLE cells, an RNeasy Mini Kit^®^ (Cat. No. 74104, Qiagen) was used for RNA purification, according to the manufacturer’s instructions. Purified RNA underwent Bioanalyzer analysis and samples with RIN (RNA Integrity Number) values greater than 9.0 were selected. These RNA samples were submitted to the JP Sulzberger Columbia Genome Center, New York, USA. RNA underwent Poly-A purification to enrich mRNAs, and library construction was done using Illumina TruSeq chemistry. Libraries were sequenced using an Illumina NovaSeq 6000. Samples were multiplexed in each lane, which yields a targeted number of paired-end 100bp reads for each sample. About 20 million reads were obtained for each sample.

### Cell infiltration assays

A cell migration/chemotaxis assay kit (24-well, 5µm) was purchased from Abcam (Cat. No. ab235696). HMLE-Cas9 and HMLE-FLNBΔH cells were cultured in serum-free MEGM^TM^ media for 24 hours, and 3.0x10^5^ cells were seeded in the top chamber. In the bottom chamber, serum-free media, serum-containing media or inducer-containing media were added to each well. The top chamber was placed on the bottom chamber, and the plate was incubated at 37°C in a 5% CO_2_ incubator. After 24 hours of incubation, the cells in the upper chamber were removed with a cotton swab, and the invasive cells were dyed using Cell Dye I solution included in the kit. The fluorescence created by the Cell Dye I was measured by reading fluorescence (Ex/Em = 530/590 nm). The experiment was repeated three times.

### MTT assays

5x10^3^ HMLE-Cas9, HMLE-FLNBΔH, HeLa-Cas9, HeLa-FLNAΔH and HeLa-FLNCΔH cells were seeded in 24 well plates. Cells were cultured in complete media for 5 to 7 days, and MTT assays were performed every 24 hours. For the MTT assay, media was removed, and 62.5µl of MTT solution (5mg/ml of 3-(4,5-dimethylthiazol-2-yl)-2,5- diphenyltetrazolium bromide in PBS) was added to each well. After 3 hours of incubation, the solution was carefully removed, and 250µl of DMSO was added to each well. After the violet particles were completely dissolved, the solution color density was measured at 560 nm with background at 670 nm, using a Nanodrop-UV/vis spectrometer (Cat. No. ND2000, Thermo Scientific). The experiment was repeated three times.

### Radiation and flow cytometry

Cells were seeded at a density of 1X10^5^ per 6 well culture dishes. After 24 hours, cells were treated with γ-irradiation using X-RAD 320 Biological irradiator, and grown for 72 hours at 37°C in a 5% CO_2_ incubator. Cells were then washed once with 1X PBS, trypsinized, and collected into 15ml tubes, and then centrifuged at 1000 rpm for 15 min and washed twice with cold cell staining buffer (Cat. No. 420201, BioLegend) and resuspended in 100µl of Annexin V Binding Buffer (Cat. No. 422201, BioLegend) at a concentration of 1x10^6^ cells/ml. Then, 5µl of FITC Annexin V (Biolegend Cat. No 640906) and 5µl of Propidium Iodide solution (Cat. No. P4864-10ML, Sigma-Aldrich) were added and the cells incubated for 20 min in the dark at room temperature with gentle shaking on an orbital rotator. Live/dead cell mix samples for compensation controls were obtained by killing cells in a 58°C water bath for 20 min and mixing with an equal number of live cells. This 1:1 live/dead cell mix was split into 3 tubes for unstained, FITC-only, and PI-only compensation controls. Cells were read in a BD FACS Celesta flow cytometer and 10,000 events were collected per sample. The experiment was repeated three times.

### Mitochondrial respiratory function assay

A Mitochondrial ToxGlo^TM^ Assay kit (Cat. No. G8000) was purchased from Promega. 1.0x10^4^ HeLa-Cas9, HeLa-FLNAΔH and HeLa-FLNCΔH cells were seeded in a 96 well tissue culture dish and incubated in serum- and glucose-deficient DMEM media for 24 hours. After starvation, cells were incubated in 100µl of glucose-containing or galactose-containing DMEM media for four hours. After incubation, 100µl of ATP detection reagent was added to each well for five minutes, after which luminescence from each well was measured. The experiment was repeated three times.

### Wound healing assay

Transfected A172 cells were cultured to confluence in 6 well culture dishes. Wounds were created using 200µl pipette tips. After 0, 12 or 24 hours, photographs were taken using a bright field microscope, and the remaining cell-free wound area was calculated using ImageJ. The experiment was repeated three times.

## Acknowledgements

We thank Dr William Hahn (Harvard Medical School) for the HMLE and CRISPR-edited HMLE cell lines, and Dr Tesshi Yamada (National Cancer Center Research Institute, Tokyo, Japan) for the GFP-tagged ACTN4 plasmids. We are grateful to Julia Furnari for assistance with the post-mortem brain samples. We thank Dr Raul Rabadan and Jiayu Su for discussions regarding computational analyses, and Dr Jian Zhang for suggestions during the early stages of this study. This work was supported by NIH grant R35 GM118136 to J.L.M.

