## Supplemental Figures for "Splicing dysregulation in glioblastoma alters the function of cell migration-related genes"

### Slide 1
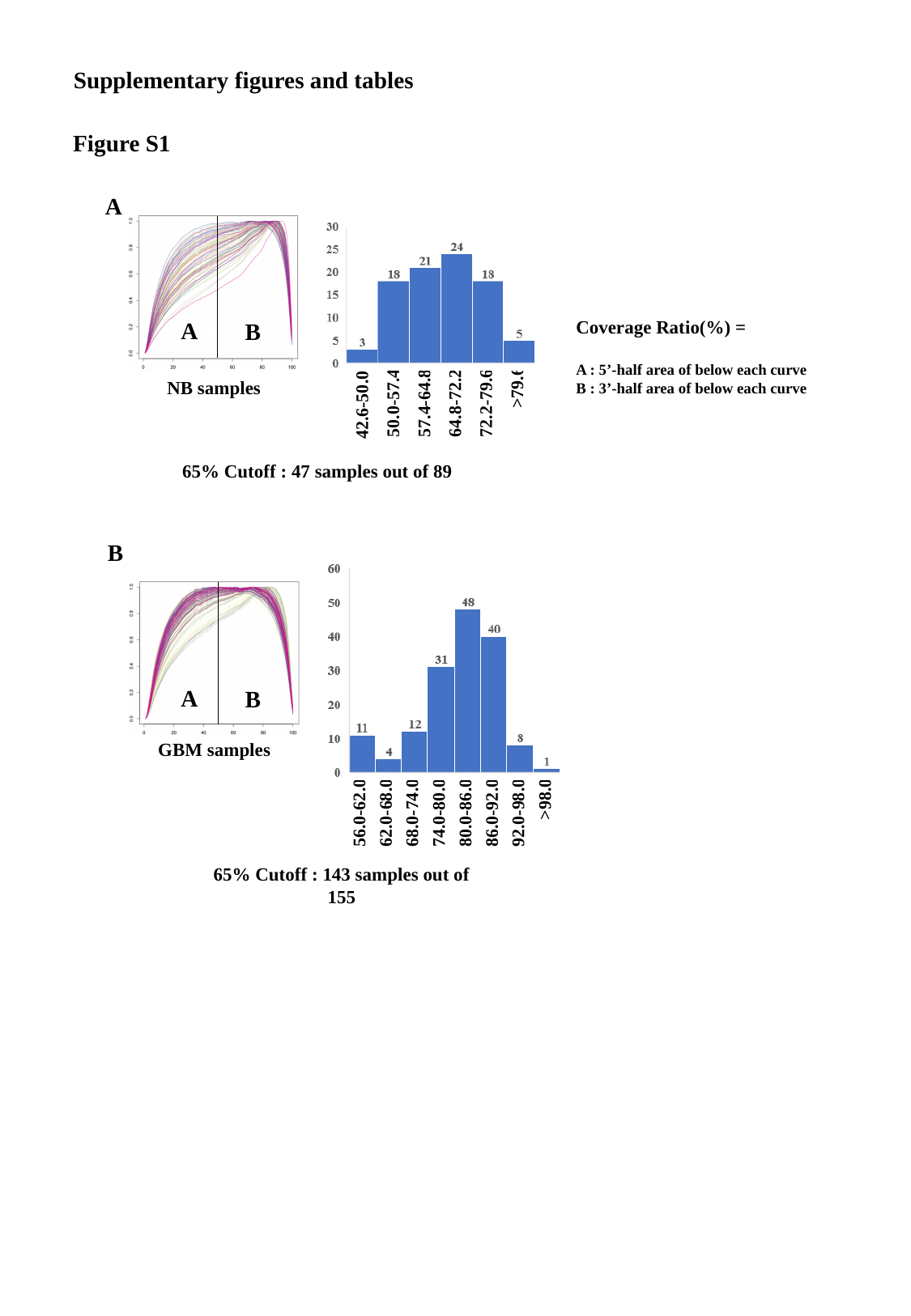

Supplementary figures and tables
Figure S1
A
A
B
NB samples
>79.6
50.0-57.4
57.4-64.8
64.8-72.2
72.2-79.6
42.6-50.0
65% Cutoff : 47 samples out of 89
B
A
B
GBM samples
>98.0
56.0-62.0
62.0-68.0
68.0-74.0
74.0-80.0
80.0-86.0
86.0-92.0
92.0-98.0
65% Cutoff : 143 samples out of 155

### Slide 2
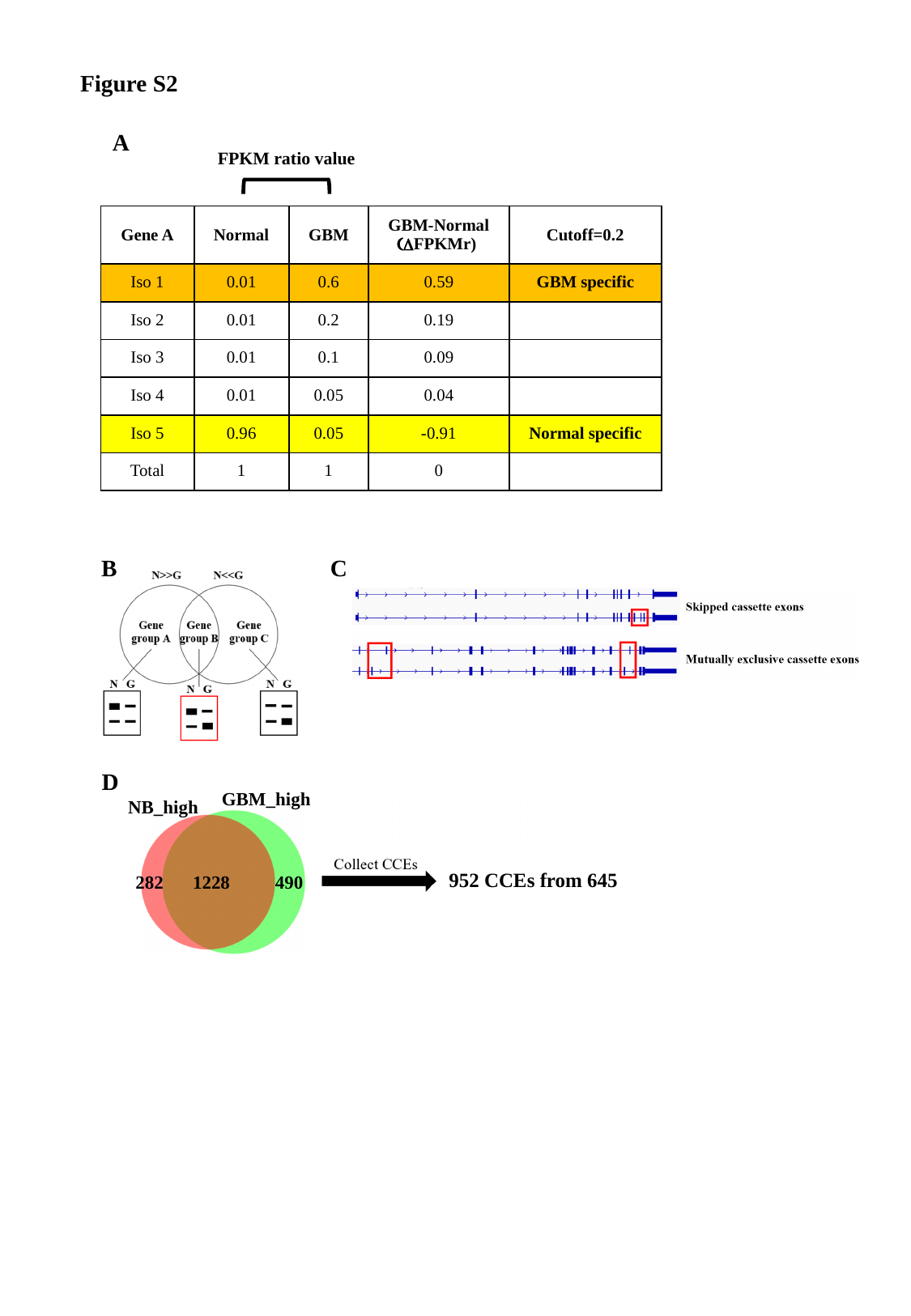

Figure S2
A
FPKM ratio value
| Gene A | Normal | GBM | GBM-Normal (ΔFPKMr) | Cutoff=0.2 |
| --- | --- | --- | --- | --- |
| Iso 1 | 0.01 | 0.6 | 0.59 | GBM specific |
| Iso 2 | 0.01 | 0.2 | 0.19 | |
| Iso 3 | 0.01 | 0.1 | 0.09 | |
| Iso 4 | 0.01 | 0.05 | 0.04 | |
| Iso 5 | 0.96 | 0.05 | -0.91 | Normal specific |
| Total | 1 | 1 | 0 | |
B
C
D
GBM_high
NB_high
282
490
1228
952 CCEs from 645

### Slide 3
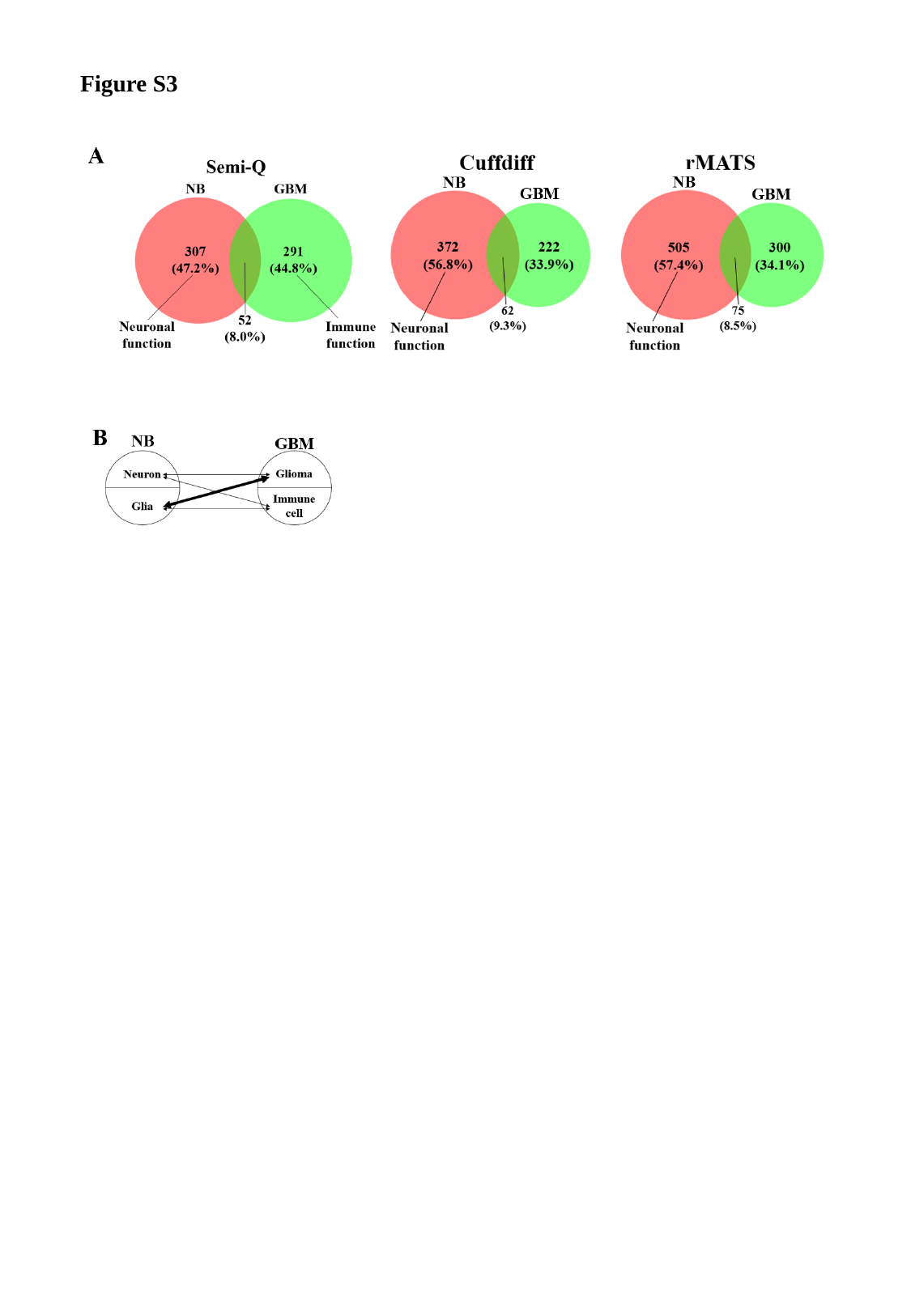

Figure S3

### Slide 4
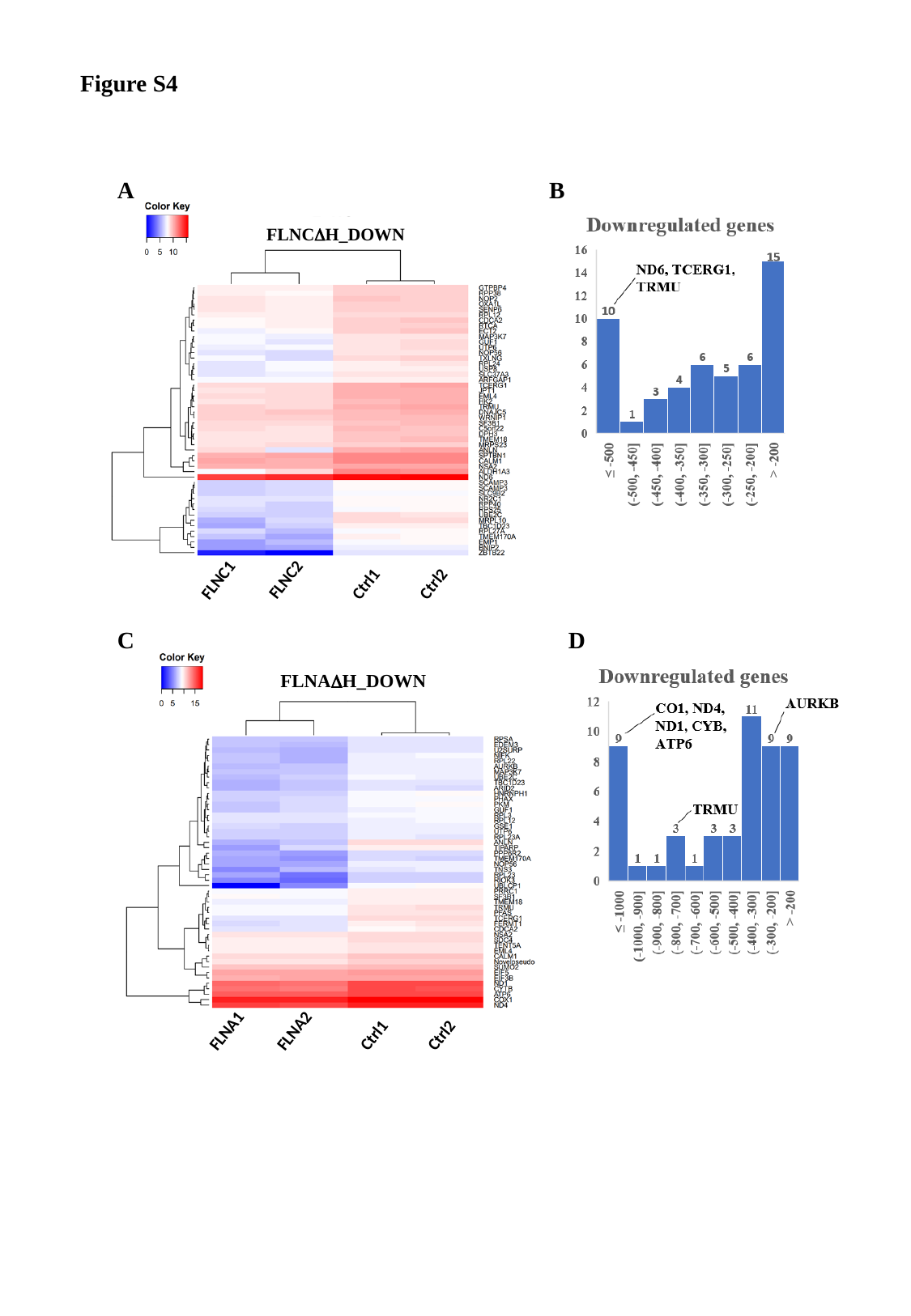

Figure S4
A
B
FLNCΔH_DOWN
FLNC2
FLNC1
Ctrl1
Ctrl2
C
D
FLNAΔH_DOWN
FLNA2
FLNA1
Ctrl1
Ctrl2

### Slide 5
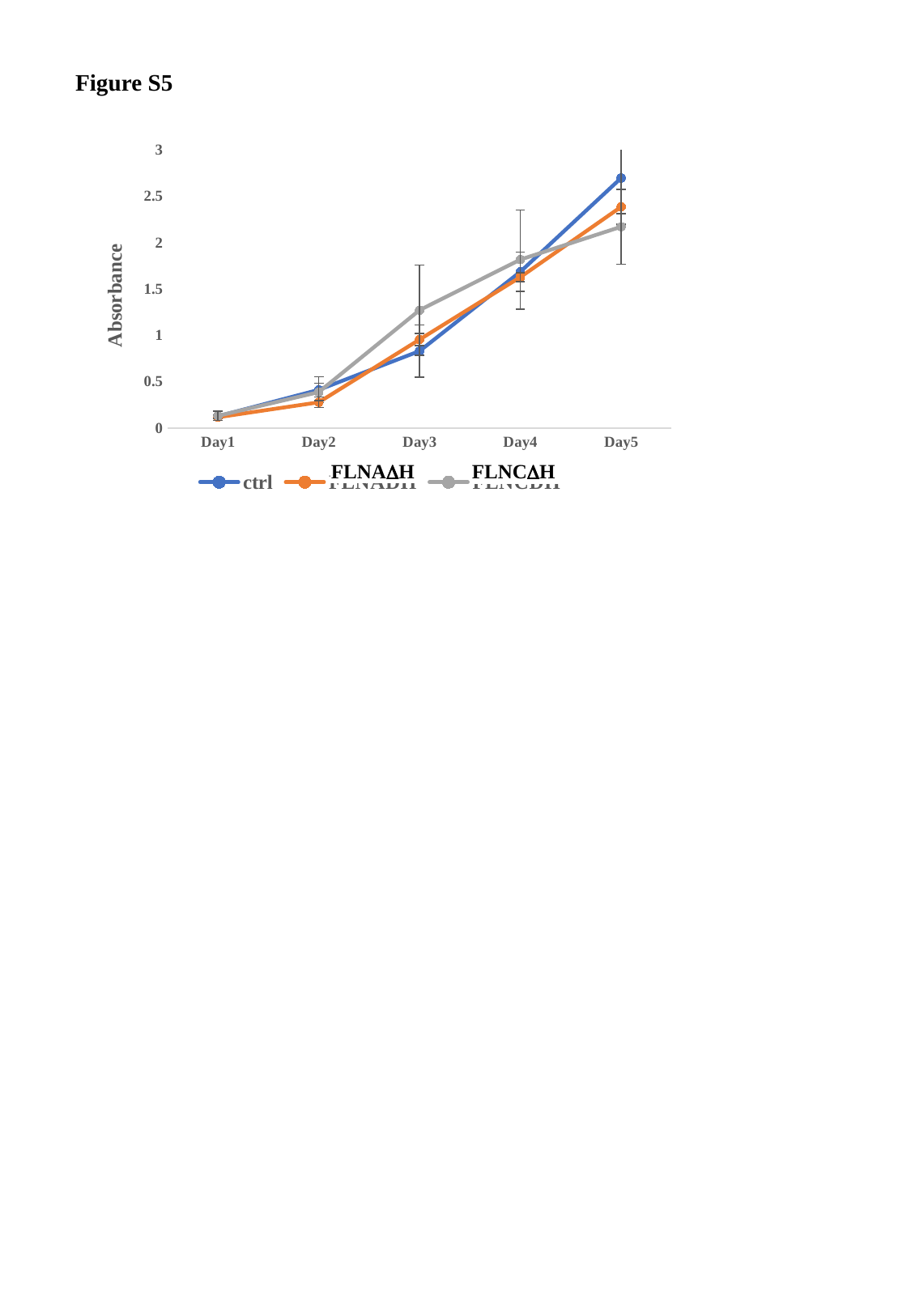

Figure S5
#### Chart
| Category | ctrl | FLNADH | FLNCDH |
|---|---|---|---|
| Day1 | 0.12733333333333333 | 0.11666666666666668 | 0.13233333333333333 |
| Day2 | 0.4123333333333334 | 0.27799999999999997 | 0.38800000000000007 |
| Day3 | 0.8303333333333334 | 0.9543333333333334 | 1.2713333333333334 |
| Day4 | 1.6849999999999998 | 1.6276666666666666 | 1.8163333333333336 |
| Day5 | 2.6956666666666664 | 2.385 | 2.170666666666667 |FLNAΔH
FLNCΔH

### Slide 6
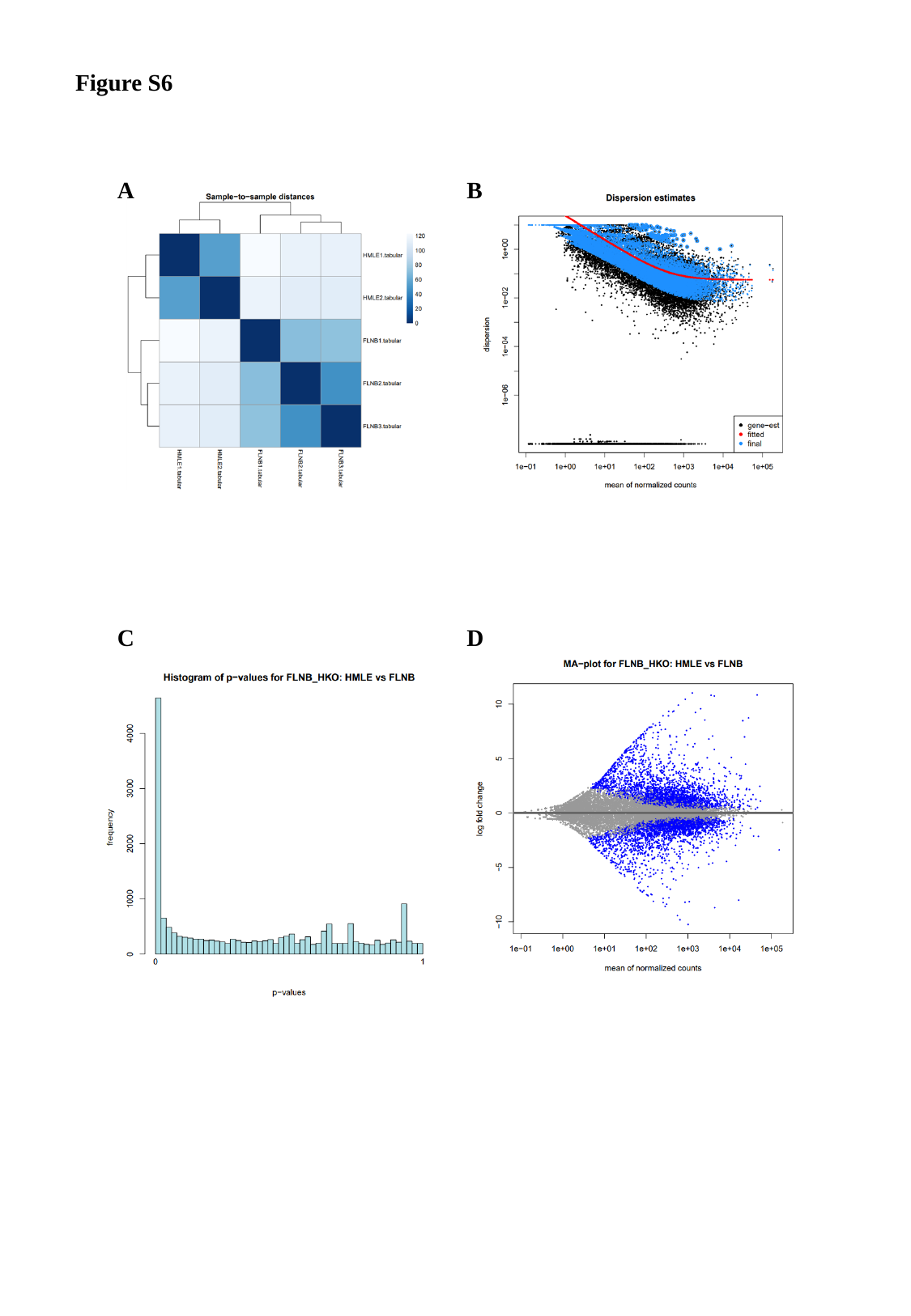

Figure S6
A
B
C
D

### Slide 7
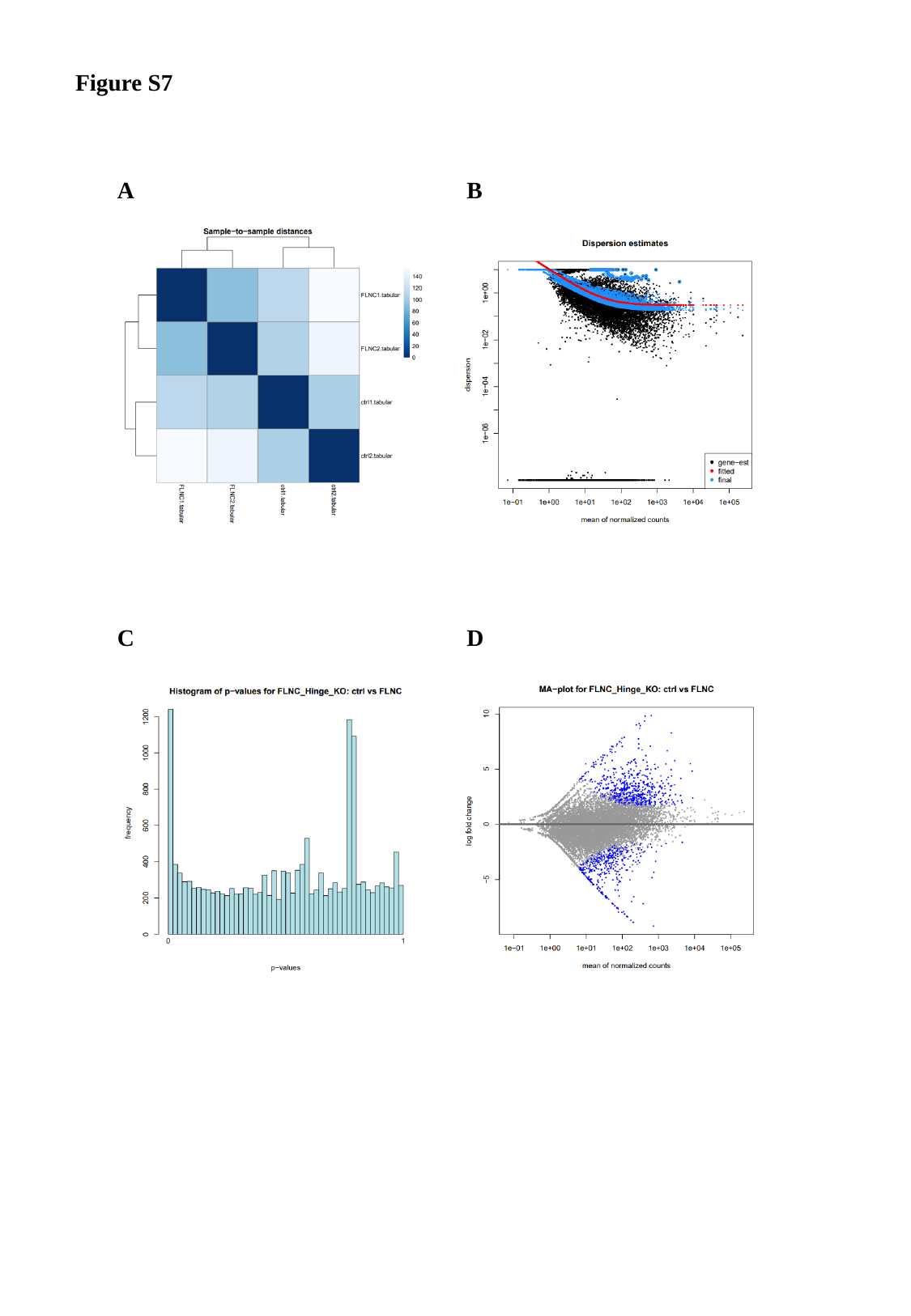

Figure S7
A
B
C
D

### Slide 8
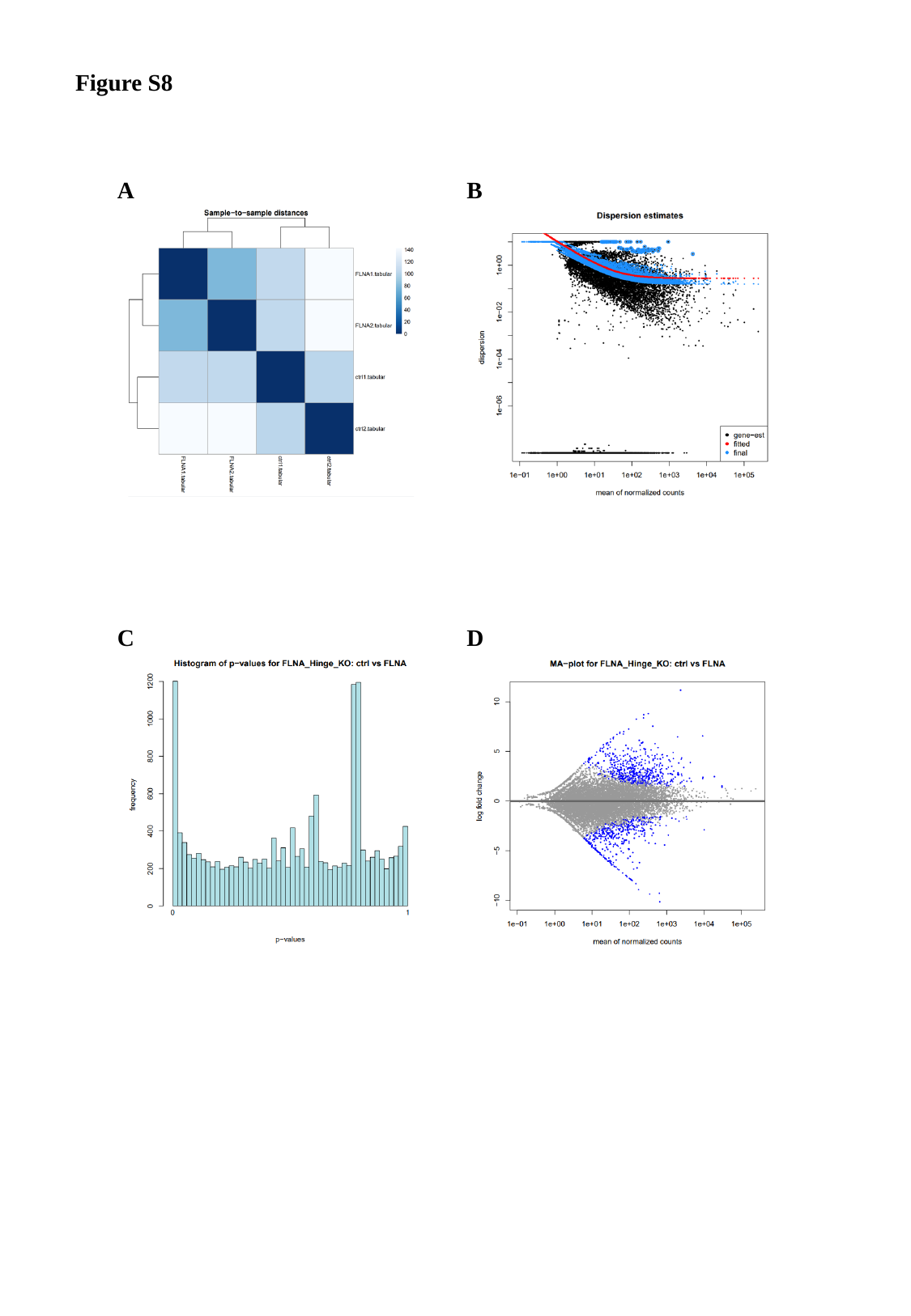

Figure S8
A
B
C
D

### Slide 9
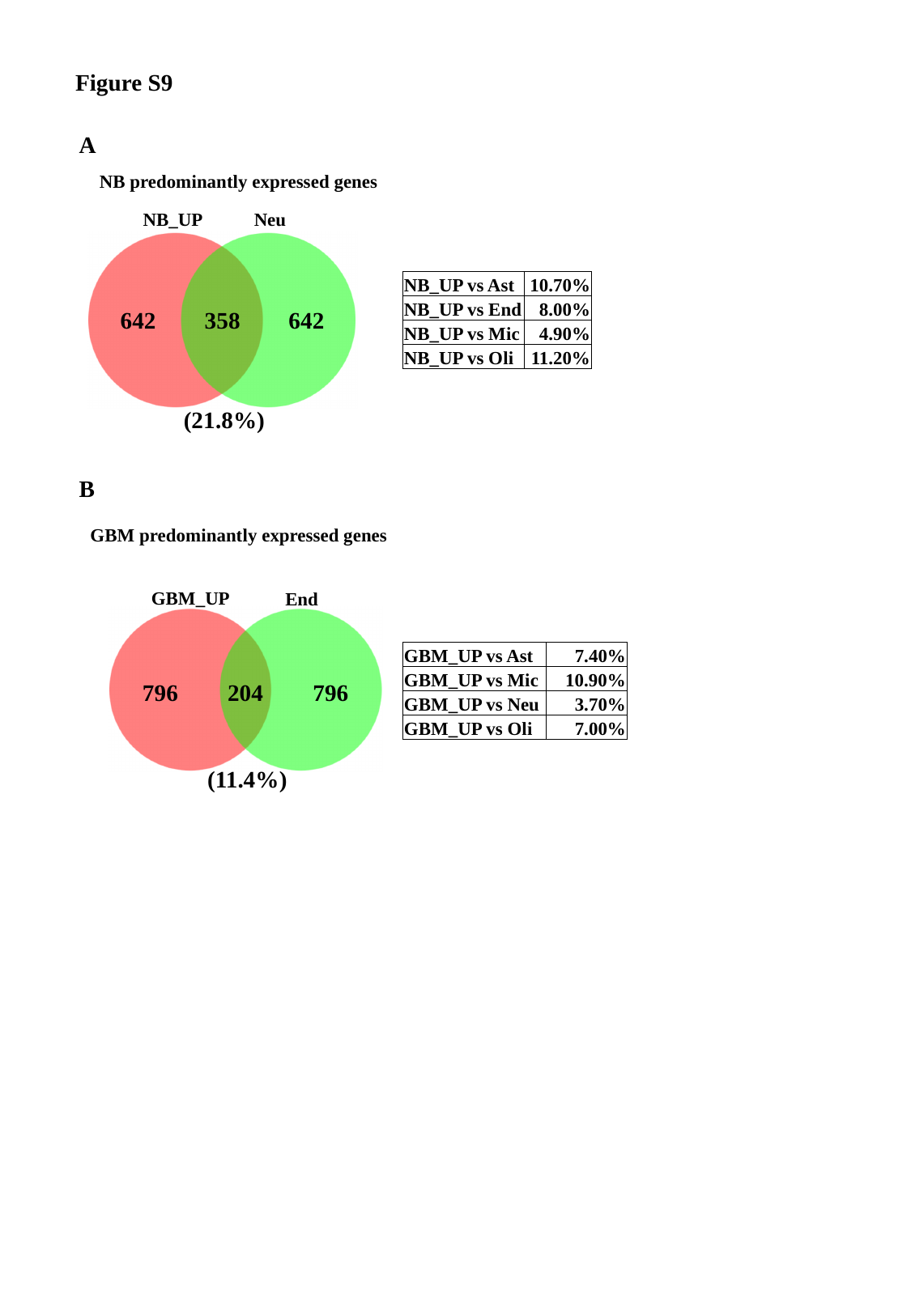

Figure S9
A
NB predominantly expressed genes
NB_UP
Neu
| NB\_UP vs Ast | 10.70% |
| --- | --- |
| NB\_UP vs End | 8.00% |
| NB\_UP vs Mic | 4.90% |
| NB\_UP vs Oli | 11.20% |
642
358
642
(21.8%)
B
GBM predominantly expressed genes
GBM_UP
End
| GBM\_UP vs Ast | 7.40% |
| --- | --- |
| GBM\_UP vs Mic | 10.90% |
| GBM\_UP vs Neu | 3.70% |
| GBM\_UP vs Oli | 7.00% |
796
204
796
(11.4%)
